# Apocarotenoid signaling regulates meristem activity and shapes shoot and root lateral organ formation in *Arabidopsis*

**DOI:** 10.1101/2024.12.09.627628

**Authors:** Julio Sierra, Lina Escobar-Tovar, Selene Napsucialy-Mendivil, Omar Oltehua-López, Joseph G. Dubrovsky, Ryan P. McQuinn, Patricia Leon

**Affiliations:** Departamento de Biología Molecular de Plantas, Instituto de Biotecnología, Universidad Nacional Autónoma de México, Cuernavaca, Morelos 62210, Mexico; Global Centre for Land-Based Innovation, Western Sydney University School of Science. Penrith, NSW 2751, Australia

**Keywords:** Meristem, root and shoot development, carotenoids, apocarotenoids, retrograde signaling, *Arabidopsis thaliana*

## Abstract

Plant carotenoids are precursors to phytohormones and signaling molecules, playing critical roles in plant development, an emerging area of research. This study investigates the function of the undefined apocarotenoid ACS1 signal in modulating plant development, particularly its impact on the morphologenesis of lateral organs and apical meristems. By modulating ACS1 levels under varying light conditions, we demonstrate its dynamic role in leaf and root development. Notably the characteristic radial leaf morphology of the *clb5* mutant reverts to normal even days post-germination, demonstrating that ACS1 is not a toxic signal but rather a key component of a biogenic retrograde signaling pathway.

Transcriptomic analysis of *clb5* seedlings at different post-germination stages underscores the critical role of ACS1 during specific developmental window. The expression profile of this mutant correlates with a proplastid stage, where even the expression of most of the genes involved in plastid biogenesis are downregulated. Furthermore, ACS1 disrupts the expression of diverse developmentally important genes, including those participating in auxin transport and signaling, leading to impaired meristem maintenance and inhibiting leaf expansion.

The effects of ACS1 extends beyond photosynthetic tissues, impacting shoot and apical root meristem organization. In particular, ACS1 affects columella cell pattering, disrupting normal gravitropic responses. These findings demonstrate that ACS1 dynamically regulates both leaf and root development, as well as meristem activity.

This study provides new insights into the role of *cis*-carotenoids as retrograde signals, functioning very early in the plastid differentiation and emphasizes the significance of plastid retrograde signaling in plant growth and development.

## Introduction

Plant fitness and homeostasis largely depend on the proper regulation of organ development and growth, which involves continuous interplay with environmental fluctuations. This regulation results from the interaction of organelles, cellular activities and external cues. Plant organogenesis is highly flexible, allowing whole organs to modulate their growth and shape to maximize their function and adapt to the environmental changes (Efroni et al., 2010). In plants, organ development and primary growth occurs along their life cycle through the activity of two meristems: the shoot apical meristem (SAM) and the root apical meristem (RAM) (Stahl and Simon, 2010; Lee and Clark, 2015). Within these meristems, a specialized population of stem cells, known as the stem cell niche, maintains the balance between the self-renewing stem cell, proliferating meristematic and differentiating cells responsible for the development of the new organs (Aichinger et al., 2012).

Homeostasis of the SAM and RAM is tightly regulated by both internal factors (i.e hormones and nutrients) and environmental cues such as light and (a)biotic stresses (Zeng et al., 2024). In diverse species, the maintenance of the SAM is controlled by a complex regulatory network including a negative feedback loop within the WUCHEL (WUS) transcriptional factor (TF) and the CLAVATA (CLV) peptide system (Gaillochet and Lohmann, 2015). These regulators maintain a balance between cell division and differentiation by controlling the expression of key genes (Schoof et al., 2000; Carles and Fletcher, 2003). In the RAM, the stem cell niche includes the quiescent cells (QC) surrounded by multipotent stem cells that upon division contribute to the RAM proliferation domain cells that later transit to elongation and differentiation zones (Perilli et al., 2012). In the RAM, several TFs also play essential roles in cell specification and maintenance (Iyer-Pascuzzi and Benfey, 2009). The WUSCHEL-RELATED HOMEOBOX 5 (WOX5), is expressed in the stem cell population within the quiescent center (QC) controlling cell division rate and maintenance. Similar to the SAM, a CLAVATA3/EMBRYO SURROUNDING REGION (CLE)-WOX5 module regulates maintenance of the stem cell niche and the differentiation process of the root cells (Stahl et al., 2009).

The maintenance of SAM and RAM, as the development of new organs derived from them, are also regulated by hormone levels, including auxin, cytokinin and ABA (Zhao et al., 2010; Pacifici et al., 2015). Among them, auxins play a critical regulatory role in meristem homeostasis and lateral organ formation in the SAM and RAM (Wang and Jiao, 2018). This regulation is accomplished upon establishing an auxin maxima facilitated by, among others, the auxin efflux PINs carriers (Grieneisen et al., 2007). The establishment of an auxin gradient from the tip to the differentiation zone of the root is critical for meristem maintenance and differentiation processes (Roychoudhry and Kepinski, 2022). Proper auxin levels in the SAM serves as morphogenetic signal to coordinate the development of the leaf (Burian et al., 2022) and roots, including gravitropic responses (Takahashi et al., 2021).

The maintenance and activity of SAM and RAM is significantly regulated by environmental cues, (a)biotic and nutritional. These regulations play major roles in adaptation and developmental plasticity by modulating the architecture of shoots and roots, which is crucial for plant survival. For example, light, nutrient availability and reactive oxygen species (ROS) regulate the activity of both SAM and RAM by affecting the expression of key TFs such as WUS or AUXIN RESPONSE FACTORS (ARFs) (Zeng et al., 2024). However, deeper understanding of the diverse signals and signaling pathways involved in apical meristem maintenance and activity remains a major challenge central to overall plant morphogenesis.

Plastids, as key metabolic hubs and environmental sensors of the plant cell, represent a source of regulatory signals that modulate the maintenance of meristem organization and function (Chan et al., 2016). Moreover, plastid differentiation is dynamic within the SAM, where undifferentiated proplastids encompass the central zone of the meristem (Charuvi et al., 2012). Recent studies demonstrate that plastid biogenesis occurs in concert with organ development, for example, the transition from proplastid to chloroplast is closely coordinated with leaf development (Andriankaja et al., 2012; Charuvi et al., 2012; Loudya et al., 2024). Moreover, these studies support that correct plastid biogenesis and function are critical for the synthesis of central molecules for meristem activity and organ development.

Plastids transmit information about their developmental (biogenic) and functional (operational) status through diverse retrograde signals, adjusting the expression of thousands of nuclear genes that play diverse roles in the cell and development.

Plastid-derived retrograde signals are diverse and generated from distinct origins and in response to different stimuli that alter plastid homeostasis or biogenesis (Chan et al., 2016; Pfannschmidt et al., 2020; Richter et al., 2023). Numerous reports demonstrate that these plastid-derived retrograde signals impact various aspects of plant development, including leaf and root development (Asano et al., 2004; Andriankaja et al., 2012; Avendaño-Vazquez et al., 2014; Van Norman et al., 2014; D’Alessandro et al., 2019). However, the nature of many of these signals and their specific roles remains poorly defined.

An emerging class of key retrograde signals, derived from the carotenoid catabolism, are known as apocarotenoids (Moreno et al., 2021; Sierra et al., 2022). Apocarotenoid synthesis requires the enzymatic or non-enzymatic cleavage of carotenoids by carotenoid cleavage dioxygenases (CCDs) or ROS, respectively (McQuinn et al., 2015). Resulting apocarotenoid signals reprogram the expression of diverse nuclear genes, modulating developmental, nutritional and stress responses.

The ζ-carotene <u>d</u>e<u>s</u>aturase (*ZDS)* mutant, *CHLOROPLAST BIOGENESIS 5* (*clb5*), exhibits a very early arrest in plastid development and unique leaf developmental aberrations (Avendaño-Vazquez et al., 2014). These phenotypes result from the accumulation of an undefined apocarotenoid signal referred to as ACS1, derived from the cleavage of phytofluene and/or ζ-carotenes by the CCD4 dioxygenase.

Importantly, most developmental defects associated with ACS1 accumulation are reverted under low light (<10 μmol m^-2^ s^-1^) with concomitant accumulation of the ACS1 precursors, suggesting light-induced ROS may also be involved in ACS1 synthesis (Escobar-Tovar et al., 2021; McQuinn et al., 2023). ACS1 accumulation results in the deregulation the expression of thousands of nuclear genes, including those encoding for chlororibosomal proteins, leading to deficient chloroplast translation. Many other differentially expressed genes (DEGs) in *clb5* participate in processes not related to plastid functions. Accordingly, the *clb5* mutant displays significant alterations in the leaf morphology, not observed in other carotenoid deficient mutants (Escobar-Tovar et al., 2021). Further, a reprogramming of the SAM into a floral meristem is observed in *clb5* grown with a prolonged carbon source (McQuinn et al., 2023). While the biochemical identity of ACS1 remains under investigation, existing evidence suggest that it may function as a “bona fide” retrograde signal of physiological importance, potentially acting as a “checkpoint” of plastid differentiation status that influences specific aspects of plant development (Loudya et al., 2021; Kendrick et al., 2022).

In this study we investigate the role of ACS1 on the maintenance of SAM and RAM and their associated organ development. By utilizing different light conditions to manipulate ACS1 levels, we were able to examine the dynamics of these responses at specific stages and tissues of leaf and root development. Our data demonstrate that ACS1 is not a toxic signal but rather a key component of a biogenic retrograde signaling pathway, which regulates the normal progression of the meristem maintenance and formation of lateral organs in response to the plastid developmental status. Notably, our findings confirm that the influence of biogenic plastid signals extends beyond photosynthetic tissues, playing a significant role in root development. The accumulation of ACS1 as a result of a mutation in *ZDS* gene causes pronounced morphological root defects, particularly in the columella cells (CC), leading to disruption in gravitropic responses. These results underscore the importance of *cis*-carotenoids as signals regulating not only shoot, but also root development.

## Results

### Low light exposure as a method to analyze ACS1 role in development

Previous studies have shown that the radially symmetrical leaf morphology present in the *clb5* mutant reverts to a normal lamina under low light (LL, 5 µmol m^-2^ sec^-1^, 16:8 h light: dark photoperiod). This reversion is linked to the accumulation of the linear ACS1 precursors, phytofluene, and ζ-carotenes (Escobar-Tovar *et al*., 2021; McQuinn *et al*., 2023). This response demonstrates that LL treatment provides a means to manipulate ACS1 accumulation by reducing its ROS-mediated synthesis, which can be exploited to further explore the role of the *cis*- carotenoids-derived ACS1 signal in plant development. (Figure 1A)

**Figure 1.**
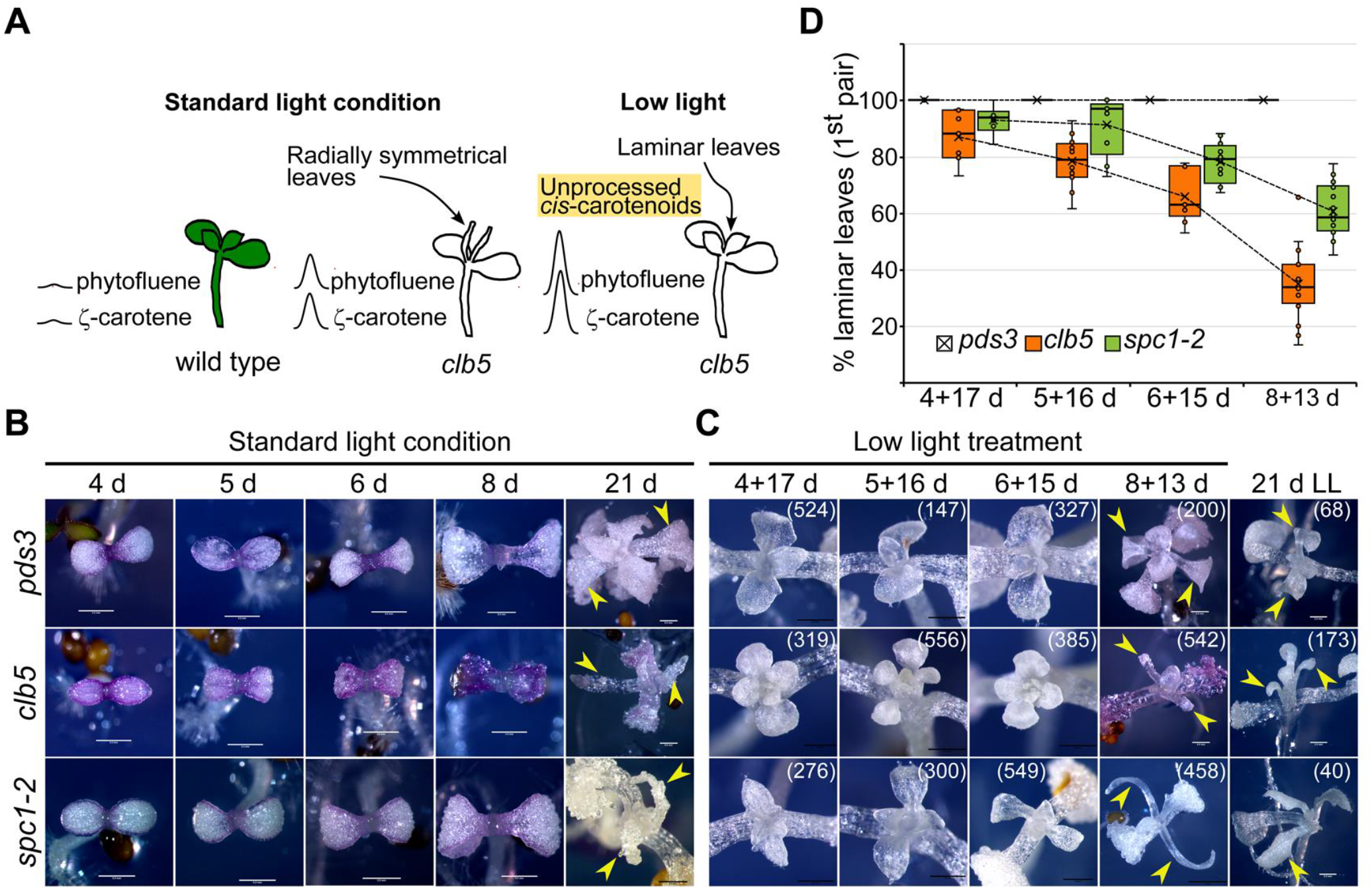
Reversion of developmental leaf defects under low light intensity at different stages of seedling development. A) Diagram of the leaf morphology under different conditions. Compared to wild-type (Wt), the *clb5* mutant displays radially symmetrical primary leaves when grown under standard light conditions (SLC) (100 µmol m^-2^ s^-1^) and accumulates low levels of the *cis*-carotenoids (phytofluene and ζ-carotenes). The leaf phenotype reverts when *clb5* is grown under low light (LL) conditions (5 µmol m^-2^ s^-1^) accompanied by a high accumulation of the ACS1 precursors, phytofluene and ζ-carotenes. B) Representative phenotypes of *pds3*, *clb5* and *spc1-2* at 4, 5, 6, 8 and 21-day-old (d) albino mutants grown in SLC. C) Representative phenotypes of 21-day-old seedlings grown in SLC or in SLC for the time indicated in each panel and transferred to LL for the indicated times (SLC+LL). The number in each panel represents the individuals analyzed across five independent experiments. D) Quantification of the laminar phenotype in the first pair of leaves in 21-day-old (d) seedlings grown in the indicated SLC+LL treatment shown in panel C. Scale bar in B and C = 0.5 mm. Yellow arrows in B and C indicate the first pair of leaves.

First, the potential dependence of the LL leaf recovery (Escobar-Tovar et al., 2021) on a specific developmental stage or growing conditions was investigated.

We examined the leaf morphology in 21-day-old *clb5* seedlings and its Col *spc1-2* allele when transferred from <u>s</u>tandard light <u>c</u>ondition (SLC, 100 µmol m^-2^ s^-^1) to LL (SLC+LL) at different times post-germination and compared them to plants grown continuously in SLC (Figure 1B). The *pds3* mutant represents an accepted albino control, given it is carotenoid-deficient and lacks ACS1-associated developmental defects, due to inhibition of carotenogenesis upstream of ZDS (Escobar-Tovar et al., 2021; McQuinn et al., 2023).

Under SLC 85% to 90% of the 21-day-old *clb5* or *spc1-2* primary leaves exhibited radial morphology, while 100% of *pds3* displayed laminar leaves (Figure 1B).

Notably, when *clb5* and *spc1-2* mutants were transferred to LL after four or five days in SLC, the percentage (85% for *clb5* and 90% for *spc1-2*) of laminar leaf development was similar to that of both mutants grown continuously under LL, (Figure 1C and D). However, the recovery rate dropped significantly when *clb5* and *spc1-2* mutants were transferred to LL later in development, with only 38% for *clb5* and 58% for *spc1-2* developing laminar leaves respectively, after 8 days (Figure 1 C, D), suggesting critical changes in leaf development at this stage. Interestingly, seedlings transferred to LL after 8 days in SLC, exhibited laminar morphology in the second leaf pair, although the first true leaves remained radial. Thus, this LL treatment was confirmed as an effective strategy to further investigate the ACS1’s roles across different tissues, developmental stages, and environmental conditions.

### Shoot apical meristem maintenance is altered by ACS1

Previous studies have shown that under SLC, *clb5* mutant produced only two primary radial leaves, halting further organ development. In contrast, with 3% sucrose, *clb5* SAM undergoes identity reprogramming, producing 3- 4 terminal organs with floral traits (McQuinn et al., 2023). Given SAM maintenance affects leaf development and vice versa (Qi et al., 2014), we investigated potential developmental alterations in the SAM of *clb5* and *spc1-2* mutants and their relationship to ACS1 accumulation. To analyze WUS and CLV3 expression patterns for SAM organization (Somssich et al., 2016; Shimotohno, 2022), both *proWUS*::GUS and *proCLAVATA3::CLAVATA3:GUS* (CLV3-GUS) reporters (Brand et al., 2002; Baurle and Laux, 2005) were introduced into *clb5* background and compared their expression in *clb5,* to that of wild-type (Wt) Col-0 and Col-0 norflurazon-treated (NFZ) under SLC and LL conditions, as controls. NFZ inhibits the PDS3 enzyme, blocking carotenogenesis synthesis, inducing retrograde signals due to photooxidative stress, but does not affect leaf morphology similar to the *pds3* mutant. In Wt seedlings, *WUS* expression was detected in the SAM at all the stages analyzed (Figure 2A). However, in both Wt NFZ-treated and *clb5* seedlings, WUS promoter activity was undetectable even after the emergence of the second leaf pair (10-day-old), indicating low WUS expression levels due to metabolic deficiencies, as reported for photosynthetic-deficient seedlings (Pfeiffer et al., 2016). Therefore, WUS marker did not allow the assessment of meristem status in response to ACS1 accumulation. Notably, the SAM structure in *clb5* seedlings was altered, lacking the typical dome-like shape observed in Wt and NFZ-treated plants (Figure 2A).

**Figure 2.**
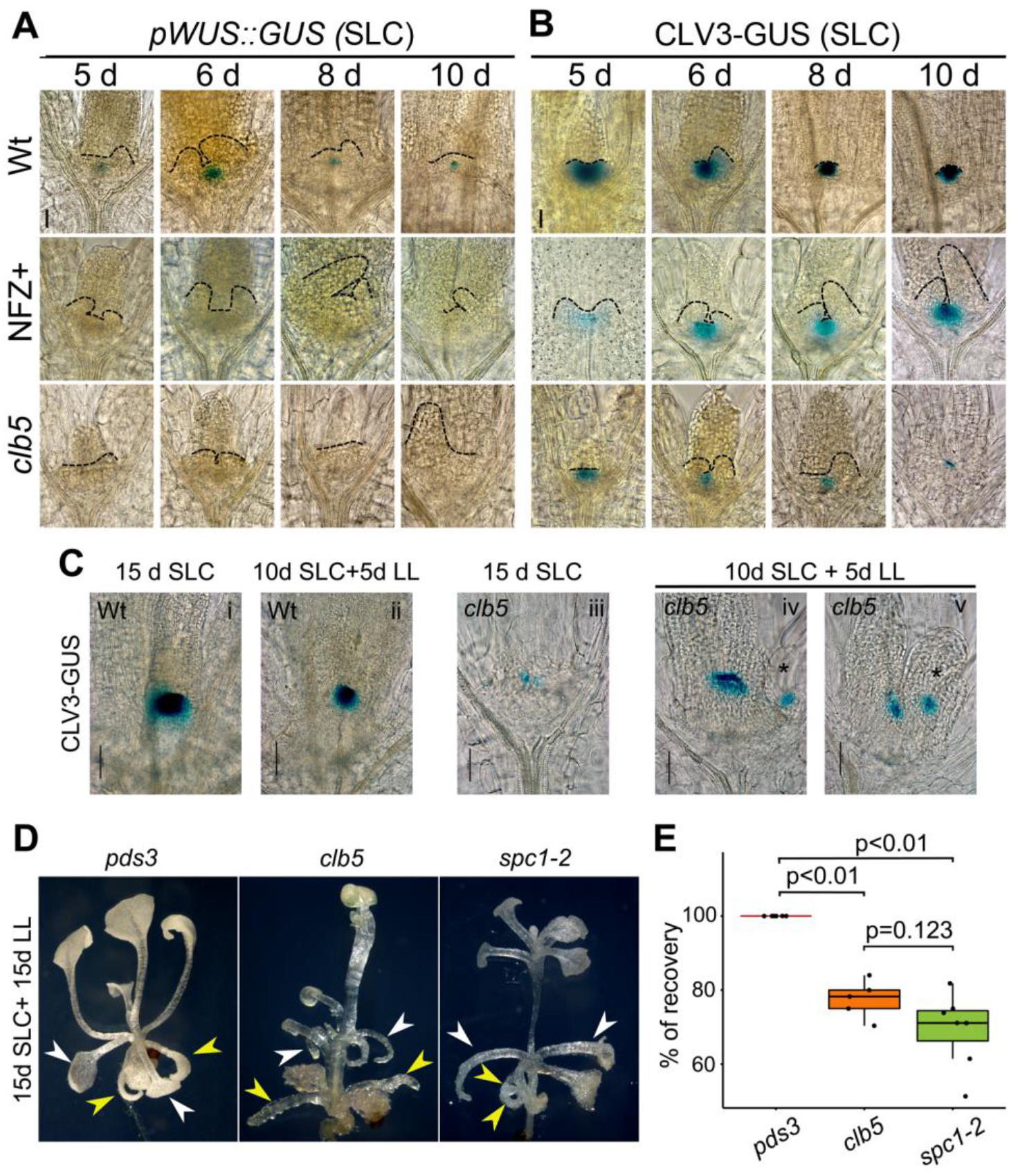
The WUS-CLV3 signaling system is affected in *clb5* causing morphological defects in the SAM that are reverted when ACS1 is reduced. Expression pattern of A) *WUSCHEL* (proWUS::GUS) and (B) *CLAVATA3* (CLV3- GUS) markers in wild-type (Wt), NFZ-treated (+NFZ) and *clb5* mutant grown in standard light conditions (SLC) during 5, 6, 8 and 10 days (d). C) CLV3 expression pattern in Wt (i) and *clb5* (iii) seedlings grown for 15 days under SLC and Wt (ii) and *clb5* (iv and v) grown 10 days in SLC plus 5 d in LL. D) Phenotypes of 30-day- old plants grown under SLC for 15 days and transferred to LL for 15 days. Arrows indicate the first (yellow) and second (white) pair of leaves. E) Percentage of laminar leaves (% recovery) shown in D in *clb5* and *spc1-2* compared to *pds3* (15SLC+15LL) across five independent analyses. Total number of pooled individuals was 451 for *pds3*, 190 for *clb5*, and 600 for *spc1-2*. Pairwise comparison was done using the Tukey HSD test. Dashed lines in A and B show the SAM architecture and developing leaves boundaries in each background. Ectopic CLV3 expression is marked with an asterisk (*) in C. Scale bar in A, B & C=50 μm. Scale bar in D= 50 mm.

Unlike WUS, CLV3 expression was clearly detected in the SAM of 5-day-old *clb5* seedlings, resembling that of Col-0-NFZ-treated seedlings and Wt plants, albeit at a reduced level (Figure 2B). While CLV3 expression increased throughout seedling development in Wt NFZ-treated plants, it decreased in *clb5,* showing especially low levels in 10-day-old seedlings (Figure 2B). This pattern suggests ACS1 signal accumulation may disrupt the WUS-CLAVATA pathway, potentially leading to the depletion of the *clb5* stem cell population.

Interestingly, when 10-day-old *clb5* seedlings with low CLV3 expression (Figure 2B) were transferred to LL for an additional five days (Figure 2Civ and v), CLV3 levels recovered significantly compared to seedlings grown in SLC for 15 days (Figure 2Ciii). This recovery highlights the plasticity of the *clb5* and *spc1-2* mutants and the dynamic effect of ACS1. Notably, 20% of these seedlings displayed ectopic CLV3 expression (Figure 2Civ and v), not observed in Wt plants under either SLC (Figure 2Ci) or after transferred to LL (Figure 2Cii). Given that CLV3 marks shoot stem cells (Brand et al., 2002), this ectopic expression likely indicates regions acquiring stem cell identity.

To further analyze *clb5*’s stem cell recovery when ACS1 levels decrease under LL, we transferred 15-day-old SLC-grown seedlings, with low CLV3 expression (Figure 4Ciii) to LL for an additional 15 days (15 LL). These 30-day-old *clb5* and *spc1-2* (15 d SLC+15 d LL) seedlings continued growing and developing new leaves with laminar phenotypes (69% and 77%), unlike those kept entirely under SLC, which arrest in development (Figures 1B, 2D and E). These results indicate that ACS1 not only alters leaf development but also fine-tune SAM activity.

### Gene expression profile in response to ACS1 accumulation highlights its critical role at specific developmental stages

Similar to the recovery analysis, meristem defects in *clb5* were not evident during early developmental stages (Figure 1 and 2B) but emerge later. To explore whether these defects correlate with specific gene expression profiles, we analyzed and compared global transcriptomic (RNA-seq) data of 8- and 18-day-old *clb5* and *pds3* mutant seedlings (Escobar-Tovar et al., 2021). These stages correspond to stage 1.0 (pre- emergence of primary leaves) and stage 1.02 (fully expanded primary leaves) as previously defined (Boyes et al., 2001), providing informative data to assess ACS1 impact in developmental transitions (Andriankaja et al., 2012; Loudya et al., 2021).

In stage 1.0 *clb5* seedlings, we identified 1479 differentially expressed genes (DEGs) (+/-1.5 log_2_FC; 673-up and 806-down) were identified compared to *pds3,* whereas the stage 1.02 *clb5* seedlings displayed over 3500 DEGs (Figure 3A and Supplementary Table S1). This supports that ACS1-mediated gene regulation is particularly critical at later stages. Although, many of the DEGs in the 1.0 *clb5* seedling stage were also present at 1.02 stage (53% upregulated and 71% downregulated genes), both DEG numbers and fold changes were higher at 1.02 stage (Figure 3A). This increase does not appear to be related to intrinsic stage differences, as no major differences in *pds3* expression profiles were observed between the two stages (Supplementary Figure S1). Notably, DEG relative expression levels (Z-scores) showed a significantly reduced expression in *clb5* compared to *pds3,* including not only PhANGs but also those involved in essential plastid biogenesis such as division, replication, import and translation (Figure 3B and Supplementary Table S2). These included the plastid-encoded polymerase (PEP) subunits, and even the NEP polymerase (RPOTp, Figure 3B) (Sriraman et al., 1998). Interestingly, the two plastid DNA polymerases (PO1A and POL1B) maintained or even increased their expression in *clb5* compared to *pds3* (Figure 3B), supporting that both genes display high expression. This expression pattern supports the idea that ACS1 halts plastid and cell development earlier than the division-to-cell expansion transition observed in the Arabidopsis *pds3* mutant (Andriankaja et al., 2012).

**Figure 3.**
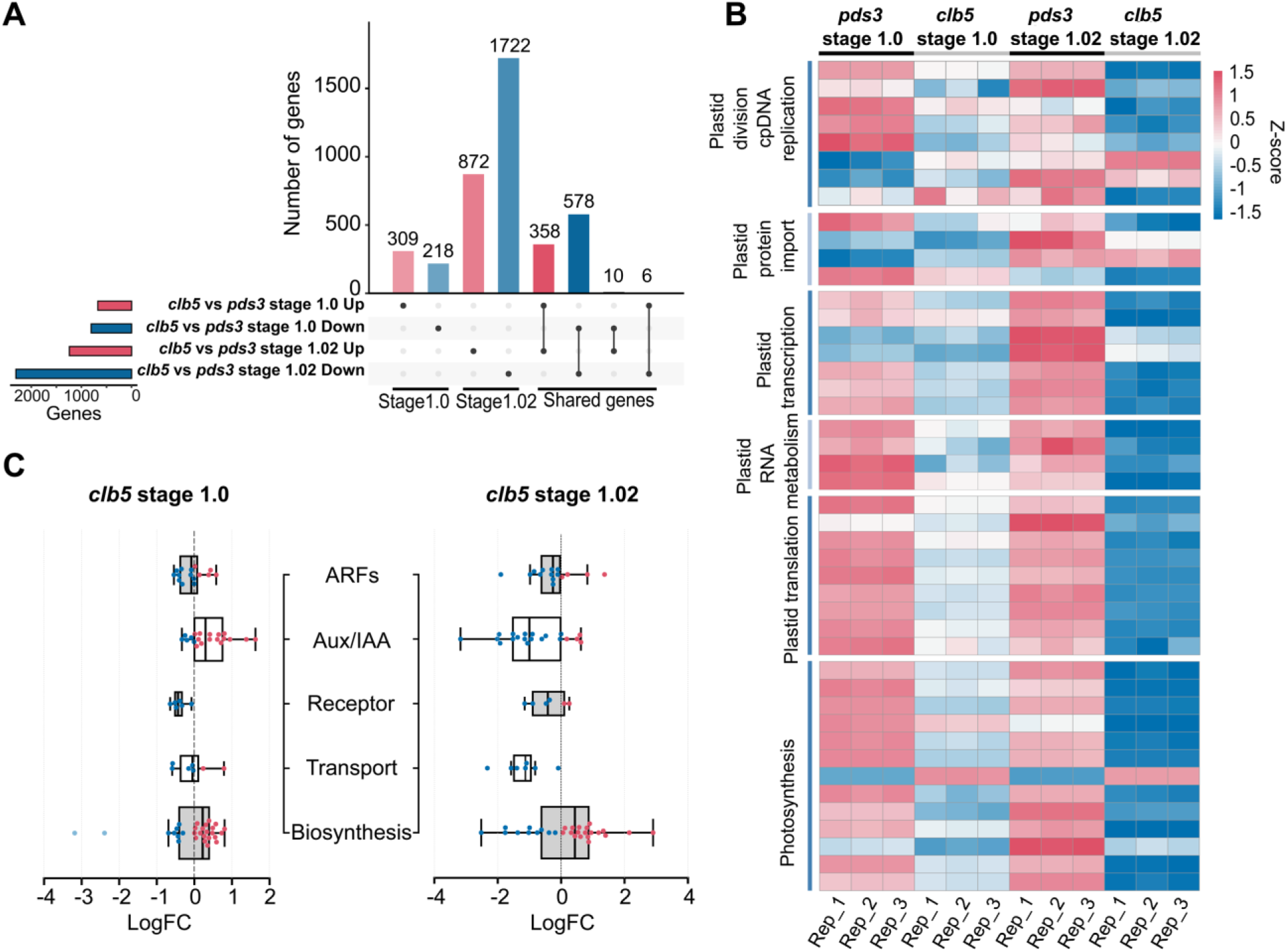
Analysis of differentially expressed genes of *clb5* along development. A) Upset plot of DEGs identified at stages 1.0 (cotyledon) and 1.02 (leaf) in the *clb5* mutant compared to *pds3* mutant, as control. B) Heatmap of genes encoding proteins located in the chloroplast of *clb5* and *pds3* using a Z- score and p-Value <0.05 of the DEGs, each condition contains three independent replicates represented in each column. C) Boxplots of expression levels of genes involved in auxin transport and signaling in *clb5* at two developmental stages; the box indicates the 25th and 75th percentile of the distribution of each gene group and the solid line indicates the median value, the T-shaped lines are the maximum and minimum LogFC values; each dot represents a gene.

**Figure 4.**
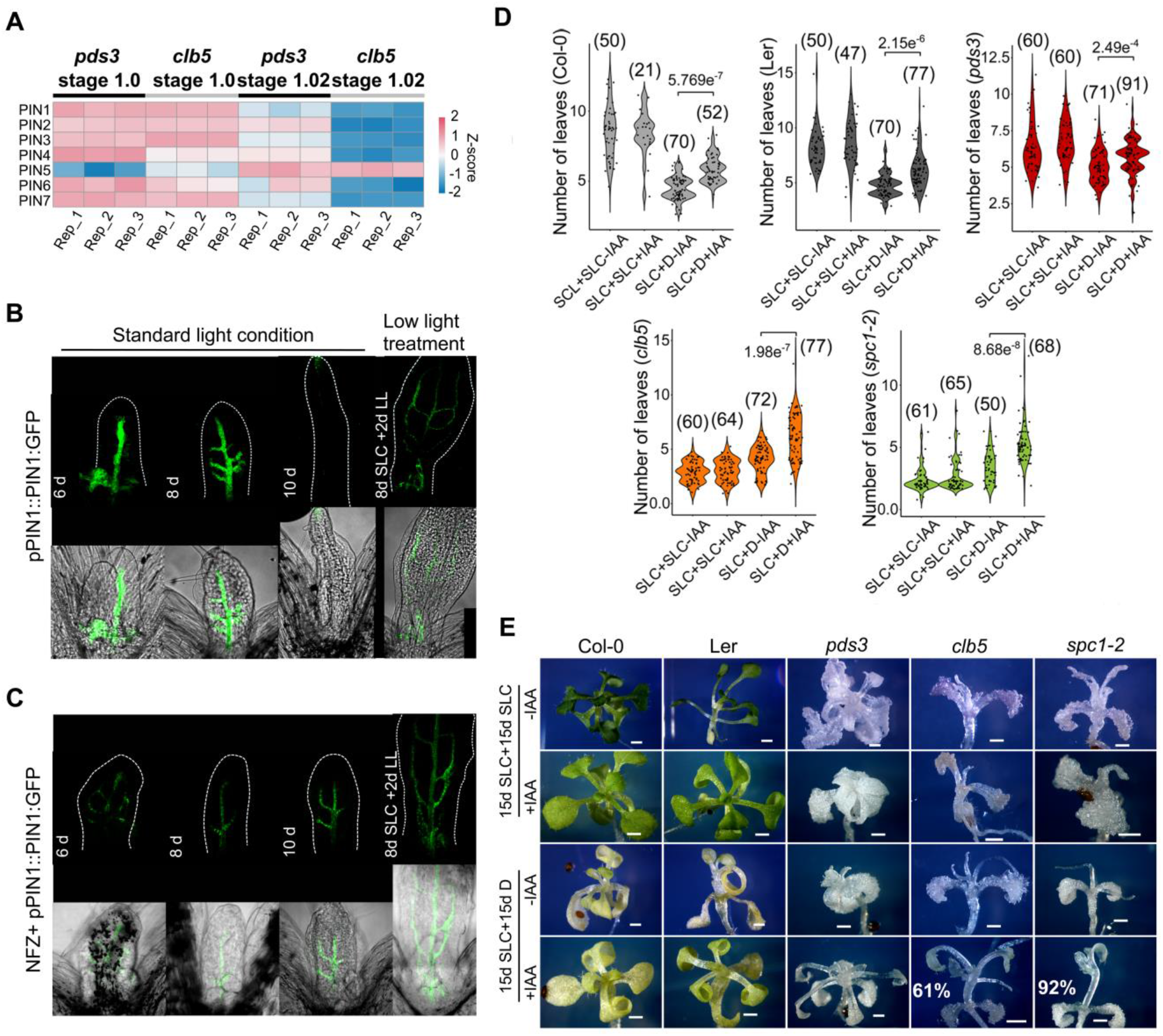
**Effect of auxins over the *clb5* and *spc1-2* leaf morphology**. (A) Heat map expression levels of genes involved in auxin transport in *pds3* and *clb5* at 1.0 and 1.02 developmental stages. Confocal images of PIN1:GFP expression in the first pair of leaves in the *clb5* mutant (B) and Wt NFZ-treated plants (C) grown in SLC or transfer to LL for two days (8d SLC+2LL). (D) Number of leaves of 30-day- old Wt (Col-0 and L*er*) and carotenoid deficient mutants *pds3*, *clb5* and *spc1-2* grown in media supplemented with 1% sucrose (+Suc) for 15-days in SLC and transferred to media with 1% sucrose and supplemented with 0.1 μM IAA (+IAA) or without (-IAA) and grow for 15 days in standard light (SLC) or darkness (D). The number in parenthesis represents the pooled individuals analyzed in four independent experiments. Kruskal-Wallis was used for multiple comparison and Dunn’s test for pairwise comparison, P-values are indicated for relevant comparisons. (E) Representative phenotypes of the 30-day-old leaves quantified in (D). The % of laminar leaves in *clb5* and *spc1-2* grown in the D+IAA is included. Scale bar =1mm.

### ACS1-related radial leaf defects are linked to auxin responses

Meristem maintenance and lateral organs formation involve various hormones, with auxins playing crucial role. Proper auxin distribution between leaf primordia and SAM is critical for organogenesis (Qi et al., 2014; Somssich et al., 2016). Comparing auxin-related gene expression between 1.0 and 1.02 transcriptomes showed significant downregulation, specially at stage 1.02, of genes involved in auxin transport and signaling, including different *PIN-FORMED* (*PIN*) and *AUXIN/INDOLE-3-ACETIC ACID* (*IAA*) genes compared to *pds3* (Figure 3C). These findings suggest that disrupted auxin homeostasis may impair meristem maintenance and organogenesis in the *clb5* mutant.

Previously, ACS1 accumulation in *clb5* leaves grown under SLC disrupts auxin accumulation and alters expression of the auxin efflux carrier PIN1 (Avendaño-Vazquez et al., 2014; Escobar-Tovar et al., 2021). Herein the transcriptomic analysis confirmed downregulation of several PIN transporters at 1.02 stage (Figure 4A). Analysis of the *proPIN1:PIN1:GFP* reporter expression in *clb5* seedlings grown in SLC at various stages of leaf development (from 6 to 10- day-old) showed that, while vascular expression was present in 6- and 8-day-old *clb5* leaves (Figure 4B), it became almost undetectable with minor accumulation at the leaf tip in 10-day-old leaves, compared with Wt Col NFZ-treated leaves (Figure 4C). This correlates well with *clb5* leaf radial symmetry becoming evident due to leaf elongation without expansion (Figure 4B). Transferring 8-day-old *clb5* seedlings from SLC to LL for two days, 40% of the emerging leaves preserved laminar morphology and PIN1-GFP expression in the developing vascular tissue (Figure 4B), indistinguishable from NFZ-treated seedlings (Figure 4C).

Auxin accumulation, inferred using the *proDR5::GUS* reporter was also disrupted. Although auxin maxima were detected at the tip of the 10 day-old *clb5* leaves under SLC, similar to Wt and NFZ-treated plants, *DR5::GUS* expression was lost in the 15 day-old *clb5* leaves (Supplementary Figure S2). However, transferring 10-day-old *clb5* seedlings to LL for five days, maintained *DR5::GUS* expression (Supplementary Figure S2). These results link ACS1 accumulation to misregulation of auxin-related genes at the primary leaf emergence (stage 1.02), affecting proper auxin distribution and leaf expansion. These findings highlight ACS1’s capacity to dynamically modulate leaf development.

Attempts to rescue laminar morphology in *clb5* and *spc1-2* seedlings with exogenous IAA (indole-3-acetic acid) in 6-day-old seedlings were unsuccessful after 9 days of treatment (Supplementary Figure 3), supporting that the ACS1- associated leaf defects in these mutants do not result from low auxin levels. To test whether auxin responses could be restored when ACS1 levels are low, Wt and mutant seedlings we grown in darkness in the presence of auxins and sugars, as under these growing conditions leaves can develop in the dark (Li et al., 2017).

Fifteen-day-old Wt, *pds3, clb5* and *spc1-2* seedlings grown under SLC were transferred to media containing sucrose with 0.1 μM of IAA and grown for an additional 15 days in the dark or under SLC. As controls, media with sucrose without IAA was included. A reduction in the number of leaves was observed in all seedlings grown in the dark with sucrose, compared to those grown under SLC in the same media, indicating meristem arrest (Figure 4D) and we did not observe significant differences in leaf number or any recovery of the radial leaf morphology in *clb5* and *spc1-2* (Figure 4D and E). However, under dark conditions with sucrose and IAA (SLC+D+IAA), *clb5* and *spc1-2* mutants showed a normal auxin response, with an increase in the number of leaves, demonstrating SAM activation (Figure 4D). More importantly, in the *clb5* (61%) and *spc1-2* (92%) new emerging leaves display laminar morphology not observed in the control without sucrose (Figure 4E and Supplementary Figure S4). These findings demonstrate that ACS1-associated leaf defects can be reversed when the ACS1 levels decline and support that *clb5* and *spc1-2* mutants retain auxin responsiveness when ACS1 levels decline.

### ACS1 influences root apical meristem length independently of auxin accumulation

Based on previous studies that show altered PIN1-GFP expression and auxin-related genes defects in *clb5* roots (Escobar-Tovar et al., 2021), we explored the potential for ACS1 to influence root development. Under SLC, primary root growth rates in *clb5*, *spc1-2* and *pds3* were lower than in their respective Wt plants (Ler and Col-0), resulting in shorter and narrower roots (Figure 5, Supplementary Figure S5). These differences were not linked to ACS1 accumulation, as similar defects occurred in *pds3*. Analysis of 10- and 15-day-old *clb5*, *spc1-2* and *pds3* roots showed shorter RAM lengths compared to Wt or NFZ-treated plants.

**Figure 5.**
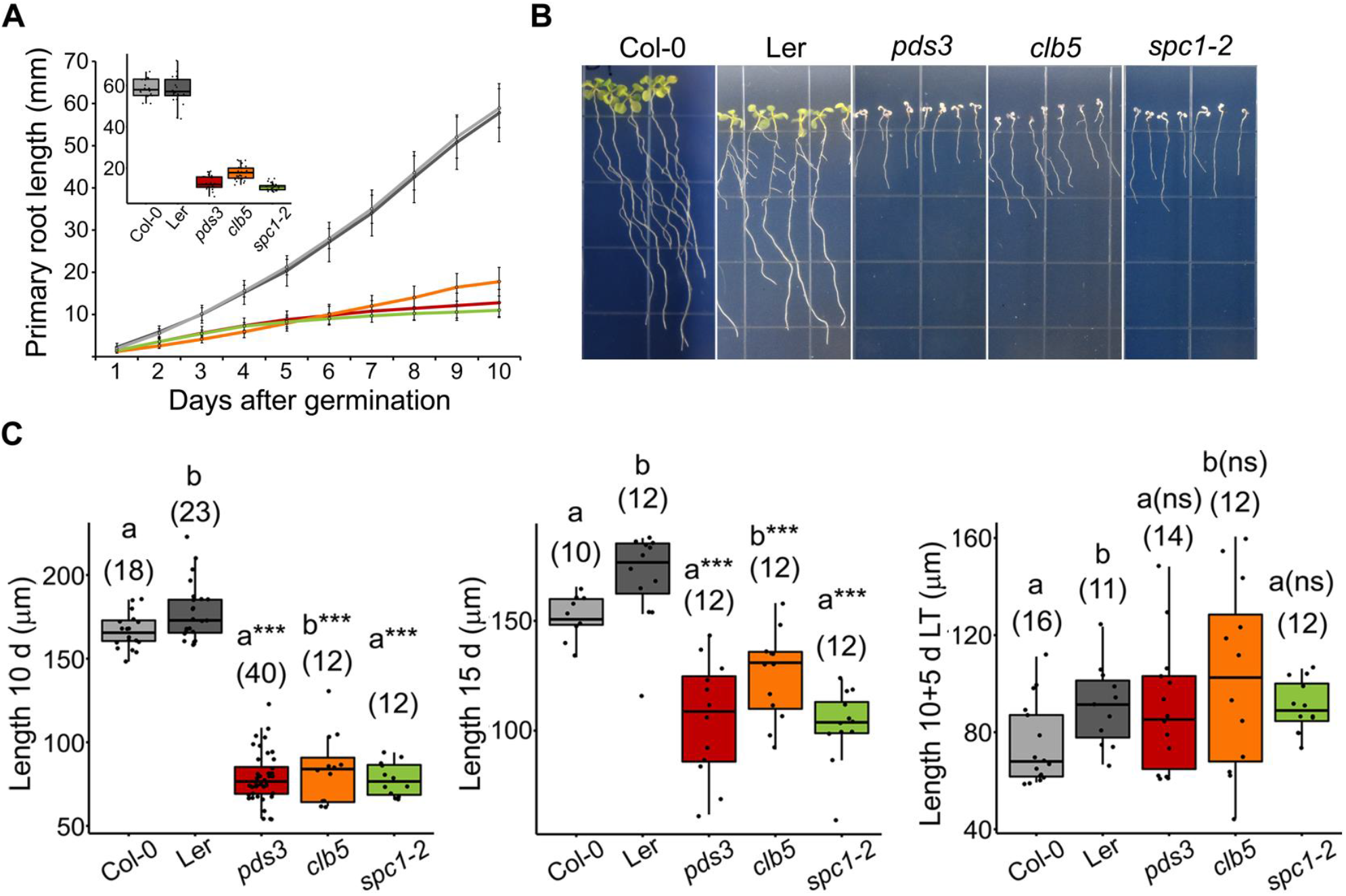
The Primary root and the RAM sizes are affected in *clb5* mutant. A) Primary root growth kinetics of Col-0 and Ler wild-type, *pds3*, *clb5* and *spc1-2* mutants analyzed along 12 days from germination. Inset shows the primary root length of the 10-day-old seedlings. Total number of pooled individuals across two independent analyses is as follows: 20 for Col-0, 20 for Ler, 22 for *pds3*, 26 for *clb5*, and 25 for *spc1-2*. B) Representative morphology of the root system of 10- day-old *pds3*, *clb5* and *spc1-2* mutants compared with the Col and Ler Wt. Quantification of the RAM length (C) in Wt (Col-0 and Ler), *pds3*, *clb5-1* and *spc1- 2* seedlings grown in SLC during 10 days (10 D) or 15 days (15 D) and grown in SLC for 10 days and 5 days in LL (10+5 D). Asterisks indicate statistically significant differences with respect to their respective Wt analyzed by Kruskal- Wallis test (P<0.01) and Dunn’s test for pairwise comparisons, lowercase letters in parenthesis indicate pairwise comparisons; ns, indicates no statistical significance. The numbers in parentheses indicate the total pooled individuals across two independent analyses.

However, these differences are not observed after five additional days of growth in LL where the length of the wild-type controls is similar to the albino mutants (Figure 5C). Notaby, auxin response, based in *DR5::GUS* expression, was similar in *clb5,* Ler and Ler-NFZ-treated plants in the different conditions tested (Supplementary Figure S6), suggesting an auxin-independent response.

Further analysis on 6-day-old *clb5*, *spc1-2, pds3* and Wt roots for responses to the addition of 0.5 µM ABA, 0.4 µM strigolactone (GR24), and 0.1, 0.2 and 0.4 µM auxin (IAA) over 9 days was carried out (Supplementary Figure S7).

Interestingly, *clb5* and *spc1-2* roots respond to exogenous auxin similarly to *pds3* and Wt, unlike aerial tissues (Supplementary Figure S7). Further, ABA enhances primary root growth in Wt and in *pds3* but not in *clb5* and *spc1-2,* while GR24 promoted root growth only in Wt, and slightly inhibited it in *clb5* and *spc1-2* (Supplementary Figure S7).

### Impact of ACS1 in the establishment and emergence of lateral root primordia

Next, we investigated ACS1’s role in lateral root (LR) development in *clb5* and *spc1-2* mutants compared to *pds3* and Wt seedlings. All three carotenoid-deficient mutants have significantly fewer LRs than Wt (Figure 5B). In addition, lateral root primordium (LRP) density was assessed. The *clb5* mutant had 50% lower LRP density than L*er* Wt (*P*=0.0029), while *spc1-2* did not differ from Col-0 Wt (*P*=0.88). Interestingly, *pds3* exhibited an increased LRP density compared to Col-0 (*P*=0.0022), highlighting different responses between these mutants (Figure 6A).

**Figure 6.**
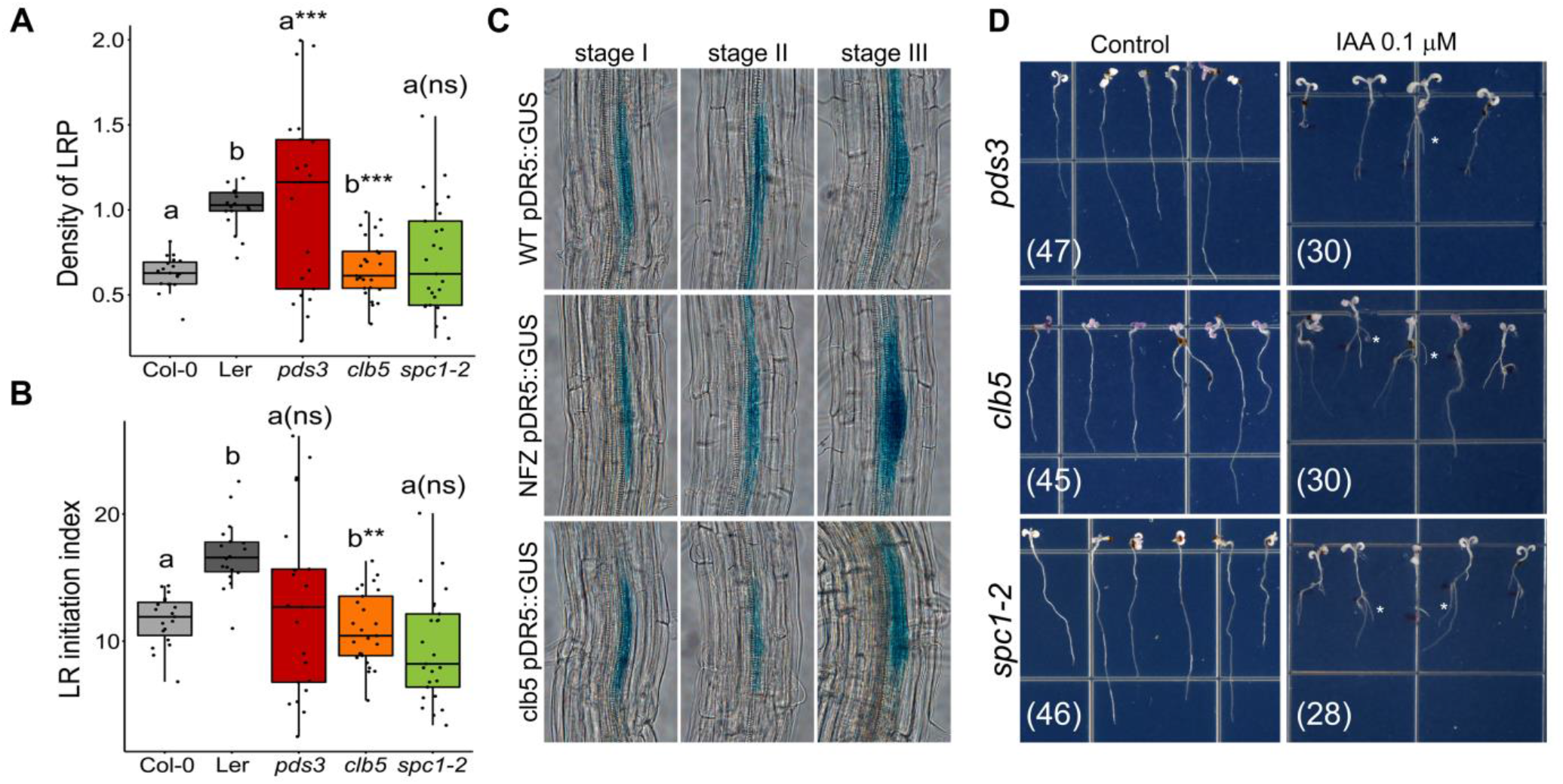
**Lateral root development is affected differently in carotenoid deficient mutants**. (A) Lateral root primordia density and (B) lateral root initiation index were measured in 12-day-old Col and Ler Wt and carotenoid deficient *pds3*, *clb5-1* and *scp1-2* mutants. Asterisks denote statistically significant differences with respect to their respective Wt (P <0.01). Letters indicate pairwise comparisons; ns indicates no statistical significance analyzed by one-way ANOVA (P <0.01) and Tukey HSD test. Total number of pooled individuals from two independent analyses is as follows: 20 for Col-0, 18 for Ler, 20 for *pds3*, 26 for *clb5*, and 25 for *spc1-2*. (C) GUS expression pattern of the *proDR5::GUS* synthetic auxin marker from Ler (WT), Ler in the presence of NFZ (NFZ) and *clb5* seedlings grown in standard conditions (SC, 100 µmol m^-2^ sec^-1^) from root initiation stages I, II and III. (D) LR emergence in *pds3*, *clb5-1* and *scp1-2* mutants in response to 0.1 μM IAA supplementation shown by the asterisks. The numbers in parentheses indicate the total pooled individuals across five independent analyses.

To evaluate LR initiation and normalize the data for cell length, we calculated the lateral root initiation Index (*l_LRI_*) (Dubrovsky et al., 2009). The *l_LRI_* was lower in *clb5* than Ler (*P*=0.00019) but similar in *spc1-2* relative to Col (*P*=0.44). This suggests that ACS1 has a clear inhibitory effect on LR initiation (Figure 6B).

Further, *pDR5::GUS* analysis showed similar auxin maxima during LRP initiation and development in *clb5* primordia to that of Wt and NFZ-treated plants (Figure 6C). These results indicate that while LR emergence defects are common in all carotenoid-deficient mutants analyzed, ACS1 accumulation specifically inhibits LR initiation.

To test whether hormones like auxins, ABA and strigolactones could rescue LR development in *clb5* and *spc1-2* mutants, 6-day-old Wt and albino seedlings were treated with 0.1 μM IAA, 0.1 μM ABA or 0.5 μM GR24 for nine days. IAA promoted LR emergence near the root differentiation zone in all mutants (Figure 6D), indicating exogenous auxin promotes emergence of existing primordia.

However, ABA or GR24 fail to promote LR emergence, suggesting these hormones do not rescue the LR deficiency under the conditions tested (Supplementary Figure S8). Overall, while ACS1 accumulation affects LR initiation and emergence, the roots remain responsive to exogenous IAA.

### ACS1 accumulation causes defects in RAM and columella morphology

To investigate the effect of ACS1 on RAM organization, 5- and 10-day-old *clb5* and *spc1-2* seedlings grown under SLC were compared to the Wt and *pds3*. While no major differences were observed in the RAM organization of 5-day-old *clb5* and *spc1-2* mutants compared to the *pds3* and Wt plants (Supplementary Figure S9A), 10-day-old mutants displayed a series of abnormalities in the stem cell niche and the columella, compared to *pds3* and Wt (Figure 7A). Although RAM cell type identities appeared to be normal and no missing cell layers were detected, the QC cell alignment was frequently disturbed in *clb5* and *spc1-2,* resulting in a disorganization of QC cells and irregular divisions in the columella initials (Figure 7Avii and viii).

**Figure 7.**
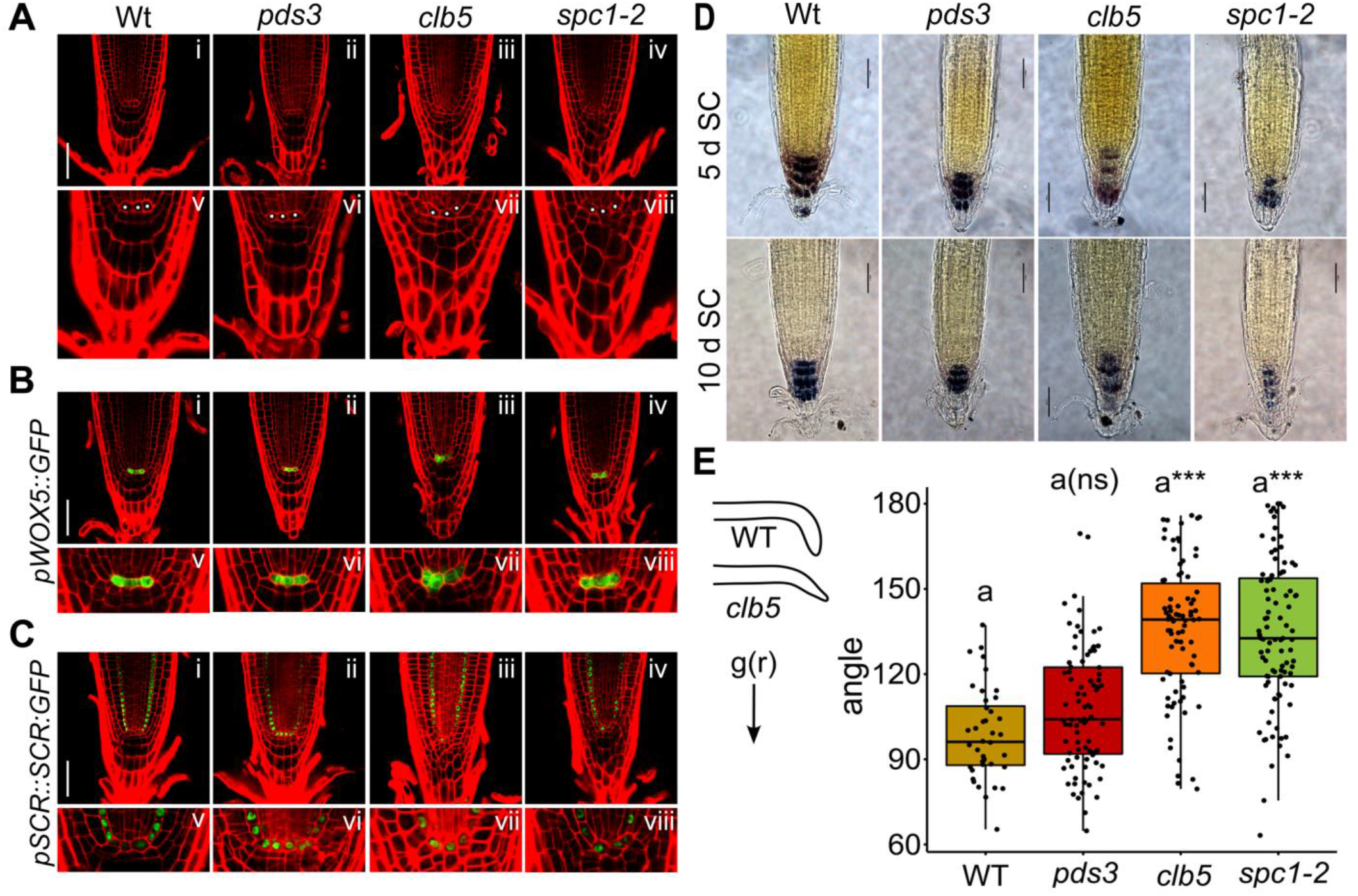
Accumulation of ACS1 affects RAM and QC structure and functionality. (A) Representative confocal images of root tip from 10-day-old morphology in wild type (Wt, A i), *pds3* (Aii), *clb5* (Aiii) and *spc1-2* (Aiv) showing the morphological defects in the *clb5-1* and *spc1-2* RAMs. Pictures below correspond to amplifications of the corresponding RAM region showing the QC and columella regions (v, vi, vii and viii). (B) Expression pattern of *proWOX5::GFP* and *proSCR::SCR:GFP* in 10-day-old Wt (Bi), *pds3* (Bii), *clb5* (Aiii) and *spc1-2* (Aiv). Pictures below correspond to amplifications of the corresponding RAM showing the QC and columella regions (v-viii). Asterisks in A correspond to the QC cells and in B to WOX5 expressing cells outside the QC line that probably result from addition divisions. Samples were stained with PI (GFP 488 nm, filter 505-525 nm) for proper visualization, The pictures are representative of three independent biological replicates n≤10. Scale bar in A=50 mm and in B and C =1 mm. (C) Starch accumulation in Wt, *pds3*, *clb5* and *spc1-2* roots analyzed by Lugol staining in 10- day-old roots (D) Boxplot of the gravitropic response of *clb5* and *spc1-2* compared to *pds3* and Wt. Inner angles were measured 48 h after 90° rotation. The diagram the gravitropic response in Wt and *clb5* mutant illustrates the observed angle in these roots. Asterisks indicate statistical significance for P<0.01 compared to their respective Wt letters indicate pairwise comparisons and statistically significant differences between genotypes was analyzed by one-way ANOVA and Tukey HSD test. ns indicates no statistical significance. Total number of pooled individuals from two independent analyses is as follows: 25 for Wt, 84 for *pds3*, 42 for *clb5*, and 56 for *spc1-2*.

To further characterize these abnormalities, we analyzed the expression of the *WOX5* and *SCARECROW (SCR)* genes which regulate RAM establishment and maintenance (Di Laurenzio et al., 1996; Haecker et al., 2004). In Wt roots, WOX5 is expressed in the QC, while *SCR* is expressed in the QC and endodermal cell lineage (Figure 7B). Confocal imaging of *proWOX::GFP* and *proSCR::SCR:YFP* markers in 5-day-old *clb5*, *spc1-2* and *pds3* mutants showed no differences compared to *pds3* and Wt (Supplementary Figure S9B and C).

However, in the 10-day-old *clb5* and *spc1-2* mutants while SCR was detected in the QC cells (Figure 7C), *WOX5* expression was also observed in misaligned QC cells (Figure 7Bviii and viii), similar to *pds3* and Wt (Figure 7 B v and vi). This indicates that while the stem cell identity appears preserved in the *clb5* and *spc1-2* QCs, ACS1 accumulation results in irregular divisions that lead to disorganization of the stem cell niche, particularly in the columella initials. This does not affect provascular and lateral-root-cap/epidermis initials cell activities.

Given the observed columella defects in the *clb5* and *spc1-2,* starch accumulation was evaluated using Lugol staining, as functional CC require amyloplasts that accumulate starch to facilitate proper gravitropic responses (Zhang et al., 2019).

Both *clb5* and *spc1-2* mutants showed a reduced starch content in CC layers at 5 and 10 days compared to *pds3* and Wt (Figure 7C). This suggests that amyloplast differentiation and starch accumulation are impaired in in *clb5* and *spc1-2*. To assess physiological impact of these changes, we analyzed gravitropic responses in 10-day-old *clb5*, *spc1-2*, *pds3* and Wt seedlings after 90° rotation. Both *clb5* and *spc1-2* displayed larger inner rotation angles (140° and 130° respectively; *P*=2.22e- 16) in comparison to Wt (90°) and *pds3* (95°; *P*=0.0099), indicating delayed gravitropic responses (Figure 7D). Overall, these results demonstrate that ACS1 accumulation in *clb5* and *spc1-2* disrupts stem cell niche organization, alters CC morphology, impairs statolith formation, and delays gravitropic responses.

### Differential expression of the ACS1 precursor enzymes

To provide insight into the regulation of ACS1accumulation within the SAM and RAM and further support ACS1’s physiological relevance, the expression patterns of key enzymes required for ACS1 precursors synthesis were analyzed (i.e. deoxy-D-xylulose 5-phosphate synthase (DXS1), PDS3 and ZDS. DXS1). DXS is part of the MEP pathway for carotenoid precursors synthesis (Cordoba et al., 2009), while PDS3 and ZDS are directly involved in ACS1 precursors synthesis (Supplementary Figure S10A) (Alagoz et al., 2018).

Single-cell transcriptomics data from the SAM showed that *DXS1*, *PSY*, *PDS3* and *ZDS* expression levels are generally low but vary within the SAM (Tian et al., 2019; Sierra et al., 2022). Specifically, *PDS3* expression was lower in the SAM central zone and higher in the rib zone, whereas *ZDS* showed the opposite pattern, suggesting regions where ACS1 could accumulate (Supplementary Figure S10B). Similar differential expression was observed in the RAM using publicly available high-resolution single-cell RNA-seq data from the Arabidopsis roots (Denyer et al., 2019), with higher accumulation of *DXS1* and *ZDS* and lower levels of *PDS3* and *PSY* in the CC (Figure 8A).

**Figure 8.**
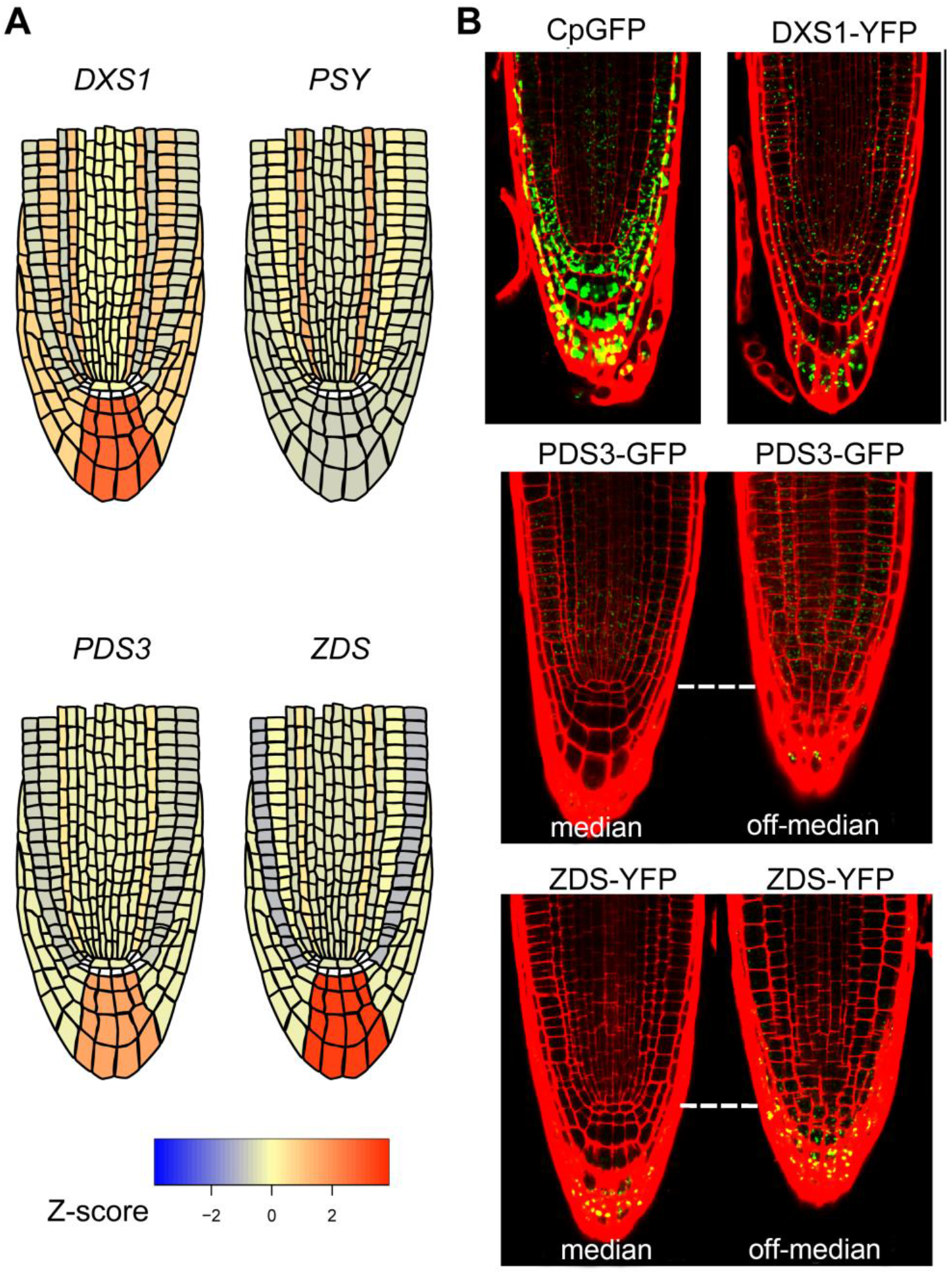
**RAM expression of carotenogenesis genes and proteins involved in ACS1 precursors**. A) Diagrammatic representation of expression levels of the carotenogenic genes. *DXS*, deoxyxylulose 5-phosphate synthase; *PSY*, phytoene synthase; *PDS3*, phytoene desaturase; and *ZDS*, ζ-carotene desaturase genes in the RAM. The color code corresponds to the expression level of the corresponding genes according to the Z-score shown for the heat map. B) Localization patterns of the plastid marker pro*35S::RBCSctp:GFP* (cpGFP), *pDXS*:DXS1-YFP, *pPDS3*:PDS3-GFP and *pZDS*:ZDS-YFP fusion proteins in the RAM of 10-day-old Wt seedlings. Pictures correspond to median (left) and off-median (right) transverse sections of the RAM. Cell membranes were stained with PI. The pictures are representative of three independent biological replicates n≥5. Barr=50 μm

To confirm the differential accumulation at the protein level, we generated transgenic lines expressing the DXS1, PDS3 and ZDS proteins from their native promoters, fused to YFP or GFP reporters. This resulted in the pro*DXS1::DXS1:YFP,* pro*PDS3::PDS3:GFP* and pro*ZDS::ZDS:YFP* lines (Supplemental Figure S11). We also included the pro*35S::RBCSctp:GFP* (cpGFP) control line for plastid localization (Wu et al., 2019). Due to the difficulty in visualizing the SAM (Supplemental Figure S10C), we focused on the RAM expression.

Consistent with the transcriptomic data, we observed low but detectable enzyme levels in the RAM. DXS1-YFP accumulates in plastids across most RAM tissues, including QC cells where the cpGFP marker was not observed (Figure 8B), indicating its crucial role throughout the RAM. In contrast, PDS3-GFP and ZDS- YFP exhibited distinct expression patterns. ZDS was more abundant in the CC and lateral root cap, but nearly undetectable in the central cylinder, endodermis and cortex (Figure 8B). Alternatively, PDS3 was present in the central cylinder, endodermis and cortex but at lower levels in the columella (Figure 8B). These findings support that different *cis*-carotenoid intermediates accumulate differentially within the RAM, creating distinct pools that serve as precursors for the synthesis of different apocarotenoids.

## Discussion

Retrograde signaling coordinates plastid functions with cellular and developmental programing, yet its impact on plant development remains poorly understood. The majority of retrograde signal identities remain unknown, and most research in this area relies on mutant analysis under various growing conditions (Cazzonelli et al., 2020).

Carotenoid-derived signals, including strigolactones, abscisic acid, and retinal, play pivotal roles in plant growth and development (Moreno et al., 2021; Sierra et al., 2022). This study expands on the role of the *cis*-carotene-derived ACS1 signal, which disrupts plastid biogenesis, causes conspicuous developmental defects, and alters the expression of hundreds of nuclear genes involved in plastid biogenesis and developmental programs. ACS1-associated leaf defects in both *clb5* or *spc1-2* mutants revert under low light conditions or in the absence of CCD4 in conjunction to the accumulation of ACS1 precursors consistent with the decrease in the ACS1 levels (Avendaño-Vazquez et al., 2014; Escobar-Tovar et al., 2021; McQuinn et al., 2023). Notably, the low-light reversion capacity persists even several days post-germination, demonstrating that ACS1 accumulation does not cause irreversible tissue damage or commit the leaf to a radial symmetry. These findings underscore the plasticity of leaf development in response to ACS1 accumulation and align with the studies emphasizing the role of retrograde signals during key leaf developmental transitions (Andriankaja et al., 2012).

Research in wheat and maize highlights the importance of biogenic retrograde signals during specific organelle differentiation stages for driving the progression of key transitions in leaf development (Loudya et al., 2021; Kendrick et al., 2022). Disruptions in chloroplast differentiation halt photosynthetic tissue development by misregulating the expression of numerous nuclear genes, a mechanism also observed in eudicots when plastid biogenesis is genetically or chemically impaired (Andriankaja et al., 2012; Loudya et al., 2020). The gene network reprogramming and morphological defects in *zds* mutants support the idea that ACS1 accumulation disrupts photosynthetic differentiation, particularly during developmental stages associated with leaf expansion.

Similar to albino monocot mutants and NF-treated-*Arabidopsis* seedlings (Kendrick et al., 2022), *clb5* mutant shows reduced expression of many PhANGs with lower levels than those in NF-treated Wt plants and the *pds3* albino mutant (Escobar-Tovar et al., 2021). However, unlike *pds3* and other albino mutants, the plastid biogenesis defects caused by ACS1 accumulation in *clb5* do not trigger compensatory gene expression responses, typically seen in *Arabidopsis* and monocots mutants (Loudya et al., 2020). Genes involved in NEP-driven plastid transcription, import, and division show significantly lower expression in *clb5* compared to *pds3*. This expression pattern resembles the *cue8 gun1* mutant in *Arabidopsis*, where plastids arrest at a very early stage of differentiation (Loudya et al., 2020). Notably, in *clb5,* the increase in the nuclear-encoded NEP (*RPOTp)* expression of the “albino compensatory syndrome” observed in *cue8 gun1* is absent, with RPOTp remaining low compared to *pds3* (Kendrick et al., 2022).

However, the organellar DNA polymerases (POL1A and POL1B) exhibit expression levels similar or slightly higher to *pds3*, indicating that ACS1 leads to a compensatory response for these particular genes. This expression profile suggests that ACS1 accumulation in *clb5* leads to a plastid biogenesis arrest at earlier developmental stage than previously reported in maize and *Arabidopsis* albino mutants, or NF-treated plants (Loudya et al., 2020; Kendrick et al., 2022). This early arrest may explain the pleiotropic defects in both leaves and roots, distinguishing *zds* from other albino mutants. Despite this, ACS1 accumulation does not cause seedling lethality, as its removal allows the continuation of plant development.

SAM and RAM serve as reservoirs of cells for organ formation, and their proper regulation is essential to balance between cell renewal and differentiation. This regulation is tightly controlled by a network of transcription factors and phytohormones. Our analysis demonstrated that in the *clb5* mutant, the WUS- CLV3 signaling system is disrupted, indicating compromised SAM homeostasis in the presence of ACS1, which hinders the development of new lateral organs.

Specifically, CLV3 transcription levels significantly decrease during *clb5* seedling development compared to Wt or Wt seedlings treated with NF. WUS and CLV3 are highly conserved regulators part of the CLV-WUS network, required for maintaining SAM homeostasis during post-embryonic development (Brand et al., 2000). The WUS-CLV3 signaling system regulates SAM activity by balancing auxin and cytokinin gradients, partially by modulating the expression of auxin and cytokinin signaling genes (Pernisova and Vernoux, 2021). This balance is critical for the proper patterning and the emergence of the leaf primordia in the SAM’s peripheral zone, where auxin accumulation promotes primordium formation (Galvan-Ampudia et al., 2020). In *clb5,* a progressive loss of *PIN1-GFP* and *proDR5::GUS* expression was also observed, correlating with disrupted auxin gradients at the SAM. This disruption aligns with the downregulation of several auxin transport and signaling genes in *clb5* mutant.

Auxin also modulates SAM homeostasis by downregulating *CLV3* expression through the AUXIN RESPONSE FACTOR 5**/**MONOPTEROS (ARF5**/**MP) TF (Luo et al., 2018). Interestingly, the transcriptomic analysis showed that ARF5 accumulates at higher levels in *clb5* compared to *pds3*, particularly at later developmental stages. This finding is intruiging and warrants future investigation.

Additionally, this study demonstrates that ACS1accumulation alters the development of the non-photosynthetic tissues, a phenomenon not reported in other carotenoid-deficient mutants, like *pds3*. These defects include shorter RAM lengths, abnormal columella development, and reduced starch accumulation in columella cells, impairing normal gravitropic responses. Accordingly, previous research has shown that plastid translation is critical for maintaining stem cell pattering in lateral root (Nakata et al., 2018). Herein, the findings further support the important function of plastid status in the root development.

Gravity sensing is important for plant nutrient absorption and environmental responses, initiated with the sedimentation of the amyloplasts within the CC (Morita, 2010). Reduced starch accumulation in amyloplasts has been linked to altered root gravitropism, highlighting the direct role of starch content in regulating this response (Sack, 1997). CCs differentiate from the columella stem cell initials, a process tightly coordinated with the biogenesis of amyloplasts from proplastids.

During differentiation, statoliths (specialized amyloplasts) in CCs, accumulate starch and develop specific morphology and functionality. However, the molecular mechanisms governing this differentiation process remain poorly understood (Morita, 2010; Chen et al., 2023).

In *clb5* and *spc1-2* mutant roots, the number of starch granules in CC appeared reduced, suggesting disrupted amyloplast differentiation. Various apocarotenoid signals and hormones influence root development. For example, ABA, strigolactones and retinal influence root architecture, while ß-cyclocitral promotes cell division in the root meristem and enhances root branching (Dickinson et al., 2019; Dickinson et al., 2021; Moreno et al., 2021). Interestingly, our findings suggest that ACS1 plays a distinct role in columella development that does not overlap with these other apocarotenoids. This is evident because the *pds- 3* mutant, which disrupts β-carotene-derived apocarotenoid synthesis, does not exhibit the altered CC morphology or gravitropic defects observed in *clb5*. Therefore, the mechanism by which ACS1 impacts the organization of the stem cell niche and columella initials remains unclear and requires further investigation.

To support optimal plant growth, continuous developmental adjustments are necessary in response to environmental and metabolic cues. By modulating the accumulation of ACS1 levels, our study highlights the pivotal role of plastid status in regulating broad aspects in plant development. This regulation impacts not only photosynthetic tissues but also other organs such as SAM and RAM maintenance. Taken together our findings, we hypothesize that a tissue- or organ-specific ACS1 synthesis may create gradients of this carotenoid-derived signal, reflecting the undifferentiated plastid status and delaying specific developmental transitions in leaves and CC. In this context, ACS1 likely plays an important role in maintenance of the meristem integrity and proper organ differentiation, with broader implications for plant growth and adaptative response mechanisms.

## Material and Methods

### Plant material and growing conditions

This work used *Arabidopsis thaliana* ecotypes Columbia-0 (Col-0) and Lansberg erecta (Ler), along with previously reported mutants *clb5* (Gutiérrez-Nava et al., 2004), *pds3-1* (Qin et al., 2007) and *spc1-2* (SALK_033039) (Dong et al., 2007). Reporter lines included *proWOX5::GFP* (Sarkar et al., 2007), *proSCR::SCR:GFP* (Heidstra et al., 2004), *proDR5::GUS* (Scarpella et al., 2010) and *proWUS::GUS* (Baurle and Laux, 2005). The *proCLV3::CLV3:GUS* was donated by Dr. S. de Folter and pCP-GFP by Dr. R. Bock (Wu et al., 2018).

Seedlings were grown on 1X GM media (Murashige and Skoog with Gamborg vitamins, Phytotechology Labs, USA) with 1% (w/v) sucrose and 0.8% (w/v) phytoagar and stratified at 4°C for 4 days. Growth conditions were 16:8 h light:dark cycle at 22°C under either 100 µmol m^-2^ s^-1^ (standard light conditions, SLC) or 5 µmol m^-2^ s^-1^ (low light LL). For LL treatments, seeds germinated under SLC were transferred to LL at specified times, and phenotypes evaluated in 21-day-old seedlings. For dark treatments 15-day-old SLC-grown seedlings were transferred to darkness for 15 days, assessing phenotypes at 30 days. Mature plants were grown in a mixture of Peat moss 3 (Sunshine, Sun Gro Horticulture, Agawam, USA), vermiculite (Sun Gro Horticulture), perlite (Agrolita, Tlalnepantla, Mexico) (5:3:2) mix with Osmocote (1.7 kg/m^3^, Everris, Geldermalsen, Netherlands) under 16:8 h light:dark photoperiod at 22°C and 65 µmol m^-2^ s^-1^.

Reporter lines *proWOX5::GFP*, *proSCR::SCR:GFP*, proPDS3::PDS3:GFP were crossed with heterozygous *clb5, spc1-2* and *pds3* mutants, while *proWUS::GUS*, *proCLV3::CLV3:GUS*, *proDR5::GUS* were crossed with *clb5*. The transgene and mutations were confirmed by the albino phenotype in the F2 generation and GUS staining. The *clb5* DR5:GUS reporter line was previously reported (Avendaño-Vazquez 2014).

### Hormone and norflurazon treatments

For auxin (IAA), ABA and strigolactone (GR24) treatments, heterozygous seeds of *pds3*, *clb5*, *spc1-2* and homozygous Col-0 and L*er* were germinated on GM media and transferred to GM or GM supplemented with 0.1 μM IAA (Sigma-Aldrich, St. Louis, USA), 0.2 μM IAA; 0.4 μM GR24 (Gomez-Roldan et al., 2008) or 0.5 μM ABA (Sigma-Aldrich) and 0.1 μM ABA 6 days after germination. Seedlings were grown SLC conditions and root phenotypes analyzed in 15-days-old seedlings. For dark hormones treatments, seedlings were grown in SLC conditions for 15 days, then transferred to GM with 0.1 μM IAA and grown in darkness for 15 days, with morphology was analyzed at 30-days-old. For NFZ treatment, plants were grown on GM with 0.5 μM NFZ (Sigma-Aldrich, St. Louis, MO, USA) at the indicated times.

### Histochemical GUS staining and sample clearing

Seedlings were fixed in 90% acetone for 2h at -20°C, then incubated in GUS staining solution (100 mM NaPO_4_ pH 7.2, 0.1 % Triton X-100, 10 mM EDTA, 5 mM Potassium Ferrocyanide (K_4_ F_3_ (CN)_6_), 5 mM Potassium Ferricyanide (K₃ Fe₃ (CN)_6_: H_2_O) for at least 20 min. Seedlings were vacuum infiltrated with GUS solution containing 1 mg/mL X-GLUC (Jefferson, 1987) and then incubated at 37°C for 5 h (*proCLV3::CLV3:GUS, proDR5::GUS)* or overnight (*proDR5::GUS*, *proWUS::GUS)*. Samples were then treated in 70% ethanol to remove chlorophyll. Samples and roots were cleared in HCl 0.24 N HCl, methanol 20% at 65°C for 1 h, neutralized with NaOH 7%, methanol 60% for 20 min, and washed with ethanol 70%. Tissue was rehydrated through a series of ethanol (40%, 20%, 10%) washes and mounted in 50% glycerol with 2% DMSO.

### Lugol staining

Five- and 10-day-old seedlings were fixed in FAE and cleared using ClearSee (10% xylitol, 15%sodium deoxycholate and 25% urea) solutions for 5 days, then stained with Lugol solution (Sigma-Aldrich, St. Louis, MO, USA) for 2 minutes and rinsed with distilled water. Samples were mounted for observation under an optical microscope.

### Morphological analyses and gravitropic response quantification

The LRP density was obtained counting the root primordia per millimeter of parent root. *l_LRI_*, was estimated as the number of LRs and LRP per parent root length corresponding to 100 cortical cells in a cell file (Dubrovsky et al., 2009). For gravitropic quantification, seedlings were grown vertically in GM media under SLC for 10 days and the positions of the root tip was recorded. After rotating the plants 90 degrees, root tip position was recorded again 48 h later. The internal angle was measured using ImageJ’s angle tool (Schneider et al., 2012).

### Light and Confocal microscopy

For light microscopy, plants were visualized under bright field using an optic microscope (Nikon SMZ1500). For Confocal microscopy roots and aerial tissues were incubated in 0.1% propidium iodide (PI) solution (Sigma-Aldrich, Saint Louis, MO., USA) for 10 min before imaging. Images were acquired with an Olympus *Fluoview* FV1000 multi-photon *confocal* laser-scanning *microscope* with a 30X objective and a Kalman filter of 4. Excitation wavelengths were: GFP 488 nm; YFP 488 nm; chlorophyll 635 nm, PI 543 nm. Emission signals were collected at GFP/YFP BA 505-525 nm; PI BA 560-660 nm, Chl BA655-755 nm. Maximum intensity projections were generated in FIJI (ImageJ software, US National Institutes of Health, Bethesda, Maryland, USA).

### Molecular cloning

To generate the proDXS1::DXS:YFP construct a 4000 bp fragment (1800 bp of the upstream sequence and the DXS1 coding region containing introns) was amplified using the CLAp1800GW and CLANSTP oligonucleotides (Supplementary Table S3). For the proPDS3:PDS3:GFP, a 5476 bp fragment (1800 bp upstream region and the *PDS3* coding sequence with introns) was amplified using PDS3BPFw and PDS3nstpRev primers (Supplementary Table S3). The proZDS::ZDS:YFP fusion, was generated by amplifying a 5592 bp fragment (2243 bp intergenic region, 5’ UTR and *ZDS* coding region with intron) using ZDSFw and ZDSRv primers (Supplementary Table S3). These fragments were cloned into pMDC204 (PDS3) or pHGY Gateway binary vectors (Invitrogene, Rockville, MD, USA) and introduced into Col-0 plants via *Agrobacterium tumefaciens* floral dip transformation (Clough and Bent, 1998). At least three homozygous transgenic lines per construct were selected and analyzed for expression.

### RNA seq analysis

For the RNA-seq at the cotyledon (1.0) stage, libraries were constructed and sequenced from 8-day-old *clb5* and *pds3* seedlings from three independent biological replicates, following the procedure reported (Escobar-Tovar et al., 2021) Reads were mapped to annotated genes in the *Arabidopsis thaliana* TAIR10 genome using STAR V2.7.10a. Differentially expressed genes (DEGs) were identified via the IDEAMEX platform (Jimenez-Jacinto et al., 2019) using edgeR, DEseq2, Limma and NOIseq, with a log_2_ fold-change threshold +/- 1.5, FDR < 0.05 and adjusted p-value < 0.05. Plots were create using R-Studio. Reads are available in the NCBI Gene Expression Omnibus (https://mpss.danforthcenter.org/web/php/pages/library_info.php?SITE=at_RNAseq 2&showAll=true).

## Data analysis

Boxplots were used to visualize data distribution with box indicating the 25^th^ and 75 ^th^ percentiles, the solid line indicate median, and whiskers making the maximum and minimum values. Parametric one-way ANOVA (α=0.01) was used for multiple group comparisons, followed by Tukey HSD test for pairwise means comparisons. Normal distribution was checked using the Shapiro-Wilk test (α=0.05). Where indicated, the non-parametric Kruskal-Wallis test (right-tailed Chi-Square α=0.05) and post-hoc Dunn’s test for pairwise comparison (α=0.005) were used. Violin plots were also used to show data distribution.

## Acknowledgements

We thank the LNMA Lab, Drs. Kenny A. Agreda, Dr. Nidia Sánchez-León, Dr. Cecilia Lara-Mondragón and Janett Rios-Gonzalez for technical assistance.

## Funding information

This work was funded by the DGAPA-UNAM IN207320, IN2089923 and Pew Charitable Trusts Innovation Fund 2023 grants and by the CONAHCYT PhD fellowship for JS and UNAM-PASPA for PL.

## Author contributions

JS: Designed and performed research, data analysis, figure preparation, wrote manuscript; LET: Designed and performed research; SNM: Performed research; OOL: Conducted bioinformatics analyses, RMQ: Designed research and edited manuscript; JGD: Analyzed data and edited manuscript; PL: Designed research, analyzed data, wrote manuscript and provided funding.

## Data availability

Data set can be found in *GEO:* <u>GSE152252</u>

## Supplementary Figures

**Supplementary Figure S1. Differential expression genes in the *clb5* mutant in two developmental stages.** The Venn diagrams show the comparisons of the DGEs in *clb5* mutant compared between the DEGs of each of the genotypes (*clb5* and *pds3* mutants) at the two development stages 1.0 and 1.02 (the cotyledon stage or 1.0 was used as reference).

**Supplementary Figure S2**. **Expression pattern of auxin response DR5 marker**. GUS expression pattern of the *proDR5::GUS* synthetic auxin marker from Ler (Wt), Ler in the presence of NFZ (NFZ) and *clb5* seedlings grown in standard light conditions (SLC, 100 µmol m^-2^ sec^-1^) for 10 days and 15 days or in SLC for 10 days and 5 day under low light (LL, 5 µmol m^-2^ sec^-1^) (10+5). Arrows point to the tip of the leaves. Scale bar corresponds to 0.2 mm.

**Supplementary Figure S3**. **Effect of auxins over the *clb5* and *spc1-2* leaf morphology**. Morphology of the 15-day-old *pds3*, *clb5* and *spc1-2* leaves grown in GM media or GM supplemented with 0.1 µM, 0.2 µM or 0.3 µM IAA. Seedlings were germinated in GM media in SLC (100 µmol m^-2^ sec^-1^) and transferred to GM media or GM+IAA after 6 days. Pictures represent representative seedlings (n=15, three biological replicates). Scale bar corresponds to 0.2 mm.

**Supplementary Figure S4. Leaf development of *clb5* and *spc1-2* mutants in response to auxins in the dark without sucrose**. Phenotypes of the 30-day-old Col-0, Ler, *pds3*, *clb5* and *spc1-2* seedlings grown media supplemented with 1% sucrose (+Suc) 15 days in standard light conditions (SLC) and then transferred to media with (+Suc) or without (-Suc) sucrose and supplemented with (+) or without (-) auxin (IAA) in SLC or in the dark (D).

**Supplementary Figure S5. Primary root morphology in ACS1-accumulating seedlings.** Quantification of the primary root (A), cortex (B), and epidermis (C) cell widths from Col-0 and Ler Wt seedlings and carotenoid deficient *pds3*, *clb5* and *spc1-2* mutants. Width was measured in the differentiated zone in 12-day-old roots. Asterisks denote statistically significant differences with respect to their respective Wt and letters indicate pairwise comparisons. Statistically analysis significant differences between genotypes, statistically analysis was done by one-way ANOVA and Tukey HSD test (p<0.01), ns, denotes no statistically significant differences. Total number of pooled individuals from two independent analyses is as follows: 20 for Col-0, 18 for Ler, 20 for *pds3*, 26 for *clb5*, and 25 for *spc1-2*. (B) Representative images of the primary root differentiation zone from Col-0 and Ler Wt seedlings and carotenoid deficient *pds3*, *clb5* and *spc1-2* mutants. The epidermis (E) and cortex (C) cells are indicated. Scale bar corresponds to 20 μm.

**Supplementary Figure S6**. **Auxin response in the primary root in ACS1- accumulating seedlings**. Auxin response *proDR5::GUS* marker of the primary root tip from L*er*, L*er* norflurazon-treated (NFZ), and *clb5* seedlings grown under SLC (100 µmol m^-2^ sec^-1^) for 5, 10 and 15 days, or 10-day-old in standard light condition (SLC) and 5 days under LL (10+5). Auxin maxima is detected in the RAM region in the root and the columella. Scale bar corresponds to 200 μm.

**Supplementary Figure S7. Response of the primary root to hormonal treatments.** Quantification of the primary root length in 15-day-old from Col-0 (B), and Ler (C), Wt seedlings and carotenoid deficient *pds3* (D), *clb5* (E) and *spc1-2* (F) mutants. Plants were treated with 0.5 µM ABA, 0.4 µM GR24, 0.1, 0.2 and 0.4 µM auxin for 9 days under SLC (100 µmol m^-2^ sec^-1^). Scale bar corresponds to 0.5 mm. Asterisks denote statistically significant differences with respect to their respective Wt, letters indicate pairwise comparisons. Multiple comparison was done by one-way non-parametric Kruskal-Wallis and Dunn’s test was used for pairwise comparison (α=0.005). The numbers in parentheses indicate the total pooled individuals across three independent analyses.

**Supplementary Figure S8**. **Effect of ABA and SL supplementation on lateral root development in ACS1-accumulating mutants.** Representative images of lateral root development of Col-0 and L*er* Wt seedlings and carotenoid-deficient *pds3*, *clb5* and *spc1-2* mutants after being transferred to GM media (A) or media supplemented with 0.1µM IAA (B), 0.5 μM GR24 (C) and 0.1 μM ABA (D) for 9 days. Seedlings were grown in SLC (100 µmol m^-2^ sec^-1^) and the number of LR was analyzed in 15-day-old plants. Scale bar corresponds to 50 μm.

**Supplementary Figure S9. Analysis of RAM and QC structure and expression QC markers.** A) Representative images of RAM morphology in 5-day-old old Wt (i), *pds3* (ii), *clb5* (iii) and *spc1-2* (iv) mutants and close-up views of the stem cell niche, including QC (v-viii). Asterisks denotate the QC cells (B) GFP expression pattern of the proWOX5::GFP QC marker in the RAM (i-iv) and QC (v-viii) of Wt, *pds3*, *clb5* and *spc1-2* mutants. (C) Expression pattern of endodermis marker SCR::YFP in the RAM (i-iv) and QC (v-viii) of Wt and albino mutants, with close-up views of the QC (v-viii). For optimal visualization of the structure, samples were stained with propidium iodine (GFP 488 nm, filter 505-525 nm). Scale bar corresponds to 50 μm.

**Supplementary Figure S10. Expression patterns in carotenogenic enzymes in the SAM.** A) Diagram of genes encoding enzymes involved in carotenogenesis. B) Expression patterns of the MEP and carotenogenic genes: *DXS1*, *PSY*, *PDS3* and *ZDS*. Color correspond to the expression level of the corresponding genes according to the C) Localization patterns of the plastid marker cpGFP, DXS1-YFP, PDS3-GFP and ZDS-YFP fusion proteins in the first pair of leaves and the SAM of 4-day-old Wt seedlings. The GFP and chlorophyll (Ch) signals, merged with brightfield (BF) channel allow visualization of the organ structure.

**Supplementary Figure S11. Expression pattern of pro*DXS1::DXS1:YFP,* pro*PDS3::PDS3:GFP* and pro*ZDS::ZDS:YFP* lines.** Representative expression pattern of the fusion YFP and GFP protein was examined in the 10-day-old primary roots of independent lines under a fluorescence microscopy.

**Supplementary Table S1.** DEGs shared between *clb5* and *pds3* at cotyledon (1.0) and 1.0 (cotyledon) and 1.02 (leaf) stages in the *clb5* mutant compared to *pds3* mutant.

**Supplementary Table S2.** Data for heatmap Z-score value

**Supplementary Table S3.** Oligonucleotides used in this study.

## References

1. Aichinger E, Kornet N, Friedrich T, Laux T (2012) Plant stem cell niches. Annu Rev Plant Biol 63: 615–636

2. Alagoz Y, Nayak P, Dhami N, Cazzonelli CI (2018) cis-carotene biosynthesis, evolution and regulation in plants: The emergence of novel signaling metabolites. Arch Biochem Biophys 654: 172–184

3. Andriankaja M, Dhondt S, De Bodt S, Vanhaeren H, Coppens F, De Milde L, Muhlenbock P, Skirycz A, Gonzalez N, Beemster GT, Inze D (2012) Exit from proliferation during leaf development in Arabidopsis thaliana: a not-so-gradual process. Dev Cell 22: 64–78

4. Asano T, Yoshioka Y, Kurei S, Sakamoto W, Machida Y, Sodmergen (2004) A mutation of the CRUMPLED LEAF gene that encodes a protein localized in the outer envelope membrane of plastids affects the pattern of cell division, cell differentiation, and plastid division in Arabidopsis. Plant J 38: 448–459

5. Avendaño-Vazquez AO, Cordoba E, Llamas E, San Roman C, Nisar N, De la Torre S, Ramos-Vega M, Gutierrez-Nava MD, Cazzonelli CI, Pogson BJ, Leon P (2014) An Uncharacterized Apocarotenoid-Derived Signal Generated in zeta-Carotene Desaturase Mutants Regulates Leaf Development and the Expression of Chloroplast and Nuclear Genes in Arabidopsis. Plant Cell 26: 2524–2537

6. Baurle I, Laux T (2005) Regulation of WUSCHEL transcription in the stem cell niche of the Arabidopsis shoot meristem. Plant Cell 17: 2271–2280

7. Boyes DC, Zayed AM, Ascenzi R, McCaskill AJ, Hoffman NE, Davis KR, Gorlach J (2001) Growth stage-based phenotypic analysis of Arabidopsis: a model for high throughput functional genomics in plants. Plant Cell 13: 1499–1510

8. Brand U, Fletcher JC, Hobe M, Meyerowitz EM, Simon R (2000) Dependence of stem cell fate in Arabidopsis on a feedback loop regulated by CLV3 activity. Science 289: 617–619

9. Brand U, Grunewald M, Hobe M, Simon R (2002) Regulation of CLV3 expression by two homeobox genes in Arabidopsis. Plant Physiol 129: 565–575

10. Burian A, Paszkiewicz G, Nguyen KT, Meda S, Raczynska-Szajgin M, Timmermans MCP (2022) Specification of leaf dorsiventrality via a prepatterned binary readout of a uniform auxin input. Nat Plants 8: 269–280

11. Carles CC, Fletcher JC (2003) Shoot apical meristem maintenance: the art of a dynamic balance. Trends Plant Sci 8: 394–401

12. Cazzonelli CI, Hou X, Alagoz Y, Rivers J, Dhami N, Lee J, Marri S, Pogson BJ (2020) A cis-carotene derived apocarotenoid regulates etioplast and chloroplast development. Elife 9

13. Chan KX, Phua SY, Crisp P, McQuinn R, Pogson BJ (2016) Learning the Languages of the Chloroplast: Retrograde Signaling and Beyond. Annu Rev Plant Biol 67: 25–53

14. Charuvi D, Kiss V, Nevo R, Shimoni E, Adam Z, Reich Z (2012) Gain and loss of photosynthetic membranes during plastid differentiation in the shoot apex of Arabidopsis. Plant Cell 24: 1143–1157

15. Chen J, Yu R, Li N, Deng Z, Zhang X, Zhao Y, Qu C, Yuan Y, Pan Z, Zhou Y, Li K, Wang J, Chen Z, Wang X, Wang X, He SN, Dong J, Deng XW, Chen H (2023) Amyloplast sedimentation repolarizes LAZYs to achieve gravity sensing in plants. Cell 186: 4788–4802 e4715

16. Clough SJ, Bent AF (1998) Floral dip: a simplified method for Agrobacterium-mediated transformation of Arabidopsis thaliana. Plant J 16: 735–743

17. Cordoba E, Salmi M, León P (2009) Unravelling the regulatory mechanisms that modulate the MEP pathway in higher plants. J Exp Bot 60: 2933–2943

18. D’Alessandro S, Mizokami Y, Legeret B, Havaux M (2019) The Apocarotenoid beta-Cyclocitric Acid Elicits Drought Tolerance in Plants. iScience 19: 461–473

19. Denyer T, Ma X, Klesen S, Scacchi E, Nieselt K, Timmermans MCP (2019) Spatiotemporal Developmental Trajectories in the Arabidopsis Root Revealed Using High-Throughput Single-Cell RNA Sequencing. Dev Cell 48: 840–852 e845

20. Di Laurenzio L, Wysocka-Diller J, Malamy JE, Pysh L, Helariutta Y, Freshour G, Hahn MG, Feldmann KA, Benfey PN (1996) The SCARECROW gene regulates an asymmetric cell division that is essential for generating the radial organization of the Arabidopsis root. Cell 86: 423–433

21. Dickinson AJ, Lehner K, Mi J, Jia KP, Mijar M, Dinneny J, Al-Babili S, Benfey PN (2019) beta-Cyclocitral is a conserved root growth regulator. Proc Natl Acad Sci U S A 116: 10563–10567

22. Dickinson AJ, Zhang J, Luciano M, Wachsman G, Sandoval E, Schnermann M, Dinneny JR, Benfey PN (2021) A plant lipocalin promotes retinal- mediated oscillatory lateral root initiation. Science 373: 1532–1536

23. Dong H, Deng Y, Mu J, Lu Q, Wang Y, Xu Y, Chu C, Chong K, Lu C, Zuo J (2007) The Arabidopsis *Spontaneous Cell Death1* gene, encoding a zeta-carotene desaturase essential for carotenoid biosynthesis, is involved in chloroplast development, photoprotection and retrograde signalling. Cell Res 17: 458–470

24. Dubrovsky JG, Soukup A, Napsucialy-Mendivil S, Jeknic Z, Ivanchenko MG (2009) The lateral root initiation index: an integrative measure of primordium formation. Ann Bot 103: 807–817

25. Efroni I, Eshed Y, Lifschitz E (2010) Morphogenesis of simple and compound leaves: a critical review. Plant Cell 22: 1019–1032

26. Escobar-Tovar L, Sierra J, Hernandez-Munoz A, McQuinn RP, Mathioni S, Cordoba E, Colas des Francs-Small C, Meyers BC, Pogson B, Leon P (2021) Deconvoluting apocarotenoid-mediated retrograde signaling networks regulating plastid translation and leaf development. Plant J 105: 1582–1599

27. Gaillochet C, Lohmann JU (2015) The never-ending story: from pluripotency to plant developmental plasticity. Development 142: 2237–2249

28. Galvan-Ampudia CS, Cerutti G, Legrand J, Brunoud G, Martin-Arevalillo R, Azais R, Bayle V, Moussu S, Wenzl C, Jaillais Y, Lohmann JU, Godin C, Vernoux T (2020) Temporal integration of auxin information for the regulation of patterning. Elife 9

29. Gomez-Roldan V, Fermas S, Brewer PB, Puech-Pagés V, Dun EA, Pillot JP, Letisse F, Matusova R, Danoun S, Portais JC, Bouwmeester H, Bécard G, Beveridge CA, Rameau C, Rochange SF (2008) Strigolactone inhibition of shoot branching. Nature 455: 189–194

30. Grieneisen VA, Xu J, Maree AF, Hogeweg P, Scheres B (2007) Auxin transport is sufficient to generate a maximum and gradient guiding root growth. Nature 449: 1008–1013

31. Gutiérrez-Nava ML, Gillmor CS, Jiménez LF, Guevara-García A, León P (2004) *CHLOROPLAST BIOGENESIS* genes act cell and noncell autonomously in early chloroplast development. Plant Physiol 135: 471–482

32. Haecker A, Gross-Hardt R, Geiges B, Sarkar A, Breuninger H, Herrmann M, Laux T (2004) Expression dynamics of WOX genes mark cell fate decisions during early embryonic patterning in Arabidopsis thaliana. Development 131: 657–668

33. Heidstra R, Welch D, Scheres B (2004) Mosaic analyses using marked activation and deletion clones dissect Arabidopsis SCARECROW action in asymmetric cell division. Genes Dev 18: 1964–1969

34. Iyer-Pascuzzi AS, Benfey PN (2009) Transcriptional networks in root cell fate specification. Biochim Biophys Acta 1789: 315–325

35. Jefferson RA (1987) Assaying chimeric genes in plants: the GUS gene fusion system. Plant Mol. Biol. Report 5: 387–405

36. Jimenez-Jacinto V, Sanchez-Flores A, Vega-Alvarado L (2019) Integrative Differential Expression Analysis for Multiple EXperiments (IDEAMEX): A Web Server Tool for Integrated RNA-Seq Data Analysis. Front Genet 10: 279

37. Kendrick R, Chotewutmontri P, Belcher S, Barkan A (2022) Correlated retrograde and developmental regulons implicate multiple retrograde signals as coordinators of chloroplast development in maize. Plant Cell 34: 4897–4919

38. Lee C, Clark SE (2015) A WUSCHEL-Independent Stem Cell Specification Pathway Is Repressed by PHB, PHV and CNA in Arabidopsis. PLoS One 10: e0126006

39. Li X, Cai W, Liu Y, Li H, Fu L, Liu Z, Xu L, Liu H, Xu T, Xiong Y (2017) Differential TOR activation and cell proliferation in Arabidopsis root and shoot apexes. Proc Natl Acad Sci U S A 114: 2765–2770

40. Loudya N, Barkan A, Lopez-Juez E (2024) Plastid retrograde signaling: A developmental perspective. Plant Cell 36: 3903–3913

41. Loudya N, Mishra P, Takahagi K, Uehara-Yamaguchi Y, Inoue K, Bogre L, Mochida K, Lopez-Juez E (2021) Cellular and transcriptomic analyses reveal two-staged chloroplast biogenesis underpinning photosynthesis build- up in the wheat leaf. Genome Biol 22: 151

42. Loudya N, Okunola T, He J, Jarvis P, Lopez-Juez E (2020) Retrograde signalling in a virescent mutant triggers an anterograde delay of chloroplast biogenesis that requires GUN1 and is essential for survival. Philos Trans R Soc Lond B Biol Sci 375: 20190400

43. Luo L, Zeng J, Wu H, Tian Z, Zhao Z (2018) A Molecular Framework for Auxin-Controlled Homeostasis of Shoot Stem Cells in Arabidopsis. Mol Plant 11: 899–913

44. McQuinn RP, Giovannoni JJ, Pogson BJ (2015) More than meets the eye: from carotenoid biosynthesis, to new insights into apocarotenoid signaling. Curr Opin Plant Biol 27: 172–179

45. McQuinn RP, Leroux J, Sierra J, Escobar-Tovar L, Frusciante S, Finnegan EJ, Diretto G, Giuliano G, Giovannoni JJ, Leon P, Pogson BJ (2023) Deregulation of zeta-carotene desaturase in Arabidopsis and tomato exposes a unique carotenoid-derived redundant regulation of floral meristem identity and function. Plant J 114: 783–804

46. Moreno JC, Mi J, Alagoz Y, Al-Babili S (2021) Plant apocarotenoids: from retrograde signaling to interspecific communication. Plant J 105: 351–375

47. Morita MT (2010) Directional gravity sensing in gravitropism. Annu Rev Plant Biol 61: 705–720

48. Nakata MT, Sato M, Wakazaki M, Sato N, Kojima K, Sekine A, Nakamura S, Shikanai T, Toyooka K, Tsukaya H, Horiguchi G (2018) Plastid translation is essential for lateral root stem-cell patterning in Arabidopsis thaliana. Biol Open 7: bio028175

49. Pacifici E, Polverari L, Sabatini S (2015) Plant hormone cross-talk: the pivot of root growth. J Exp Bot 66: 1113–1121

50. Perilli S, Di Mambro R, Sabatini S (2012) Growth and development of the root apical meristem. Curr Opin Plant Biol 15: 17–23

51. Pernisova M, Vernoux T (2021) Auxin Does the SAMba: Auxin Signaling in the Shoot Apical Meristem. Cold Spring Harb Perspect Biol 13: a039925

52. Pfannschmidt T, Terry MJ, Van Aken O, Quiros PM (2020) Retrograde signals from endosymbiotic organelles: a common control principle in eukaryotic cells. Philos Trans R Soc Lond B Biol Sci 375: 20190396

53. Pfeiffer A, Janocha D, Dong Y, Medzihradszky A, Schone S, Daum G, Suzaki T, Forner J, Langenecker T, Rempel E, Schmid M, Wirtz M, Hell R, Lohmann JU (2016) Integration of light and metabolic signals for stem cell activation at the shoot apical meristem. Elife 5

54. Qi J, Wang Y, Yu T, Cunha A, Wu B, Vernoux T, Meyerowitz E, Jiao Y (2014) Auxin depletion from leaf primordia contributes to organ patterning. Proc Natl Acad Sci U S A 111: 18769–18774

55. Qin G, Gu H, Ma L, Peng Y, Deng XW, Chen Z, Qu LJ (2007) Disruption of phytoene desaturase gene results in albino and dwarf phenotypes in Arabidopsis by impairing chlorophyll, carotenoid, and gibberellin biosynthesis. Cell Res 17: 471–482

56. Richter AS, Nagele T, Grimm B, Kaufmann K, Schroda M, Leister D, Kleine T (2023) Retrograde signaling in plants: A critical review focusing on the GUN pathway and beyond. Plant Commun 4: 100511

57. Roychoudhry S, Kepinski S (2022) Auxin in Root Development. Cold Spring Harb Perspect Biol 14

58. Sack FD (1997) Plastids and gravitropic sensing. Planta 203: S63–68

59. Sarkar AK, Luijten M, Miyashima S, Lenhard M, Hashimoto T, Nakajima K, Scheres B, Heidstra R, Laux T (2007) Conserved factors regulate signalling in Arabidopsis thaliana shoot and root stem cell organizers. Nature 446: 811–814

60. Scarpella E, Barkoulas M, Tsiantis M (2010) Control of leaf and vein development by auxin. Cold Spring Harb Perspect Biol 2: a001511

61. Schneider CA, Rasband WS, Eliceiri KW (2012) NIH Image to ImageJ: 25 years of image analysis. Nat Methods 9: 671–675

62. Schoof H, Lenhard M, Haecker A, Mayer KF, Jurgens G, Laux T (2000) The stem cell population of Arabidopsis shoot meristems in maintained by a regulatory loop between the CLAVATA and WUSCHEL genes. Cell 100: 635–644

63. Shimotohno A (2022) Illuminating the molecular mechanisms underlying shoot apical meristem homeostasis in plants. Plant Biotechnol (Tokyo) 39: 19–28

64. Sierra J, McQuinn RP, Leon P (2022) The role of carotenoids as a source of retrograde signals: impact on plant development and stress responses. J Exp Bot 73: 7139–7154

65. Somssich M, Je BI, Simon R, Jackson D (2016) CLAVATA-WUSCHEL signaling in the shoot meristem. Development 143: 3238–3248

66. Sriraman P, Silhavy D, Maliga P (1998) The phage-type PclpP-53 plastid promoter comprises sequences downstream of the transcription initiation site. Nucleic Acids Res 26: 4874–4879

67. Stahl Y, Simon R (2010) Plant primary meristems: shared functions and regulatory mechanisms. Curr Opin Plant Biol 13: 53–58

68. Stahl Y, Wink RH, Ingram GC, Simon R (2009) A signaling module controlling the stem cell niche in Arabidopsis root meristems. Curr Biol 19: 909–914

69. Takahashi K, Takahashi H, Furuichi T, Toyota M, Furutani-Seiki M, Kobayashi T, Watanabe-Takano H, Shinohara M, Numaga-Tomita T, Sakaue-Sawano A, Miyawaki A, Naruse K (2021) Gravity sensing in plant and animal cells. NPJ Microgravity 7: 2

70. Tian C, Wang Y, Yu H, He J, Wang J, Shi B, Du Q, Provart NJ, Meyerowitz EM, Jiao Y (2019) A gene expression map of shoot domains reveals regulatory mechanisms. Nat Commun 10: 141

71. Van Norman JM, Zhang J, Cazzonelli CI, Pogson BJ, Harrison PJ, Bugg TD, Chan KX, Thompson AJ, Benfey PN (2014) Periodic root branching in Arabidopsis requires synthesis of an uncharacterized carotenoid derivative. Proc Natl Acad Sci U S A 111: E1300–1309

72. Wang Y, Jiao Y (2018) Auxin and above-ground meristems. J Exp Bot 69: 147–154

73. Wu GZ, Chalvin C, Hoelscher M, Meyer EH, Wu XN, Bock R (2018) Control of Retrograde Signaling by Rapid Turnover of GENOMES UNCOUPLED1. Plant Physiol 176: 2472–2495

74. Wu GZ, Meyer EH, Richter AS, Schuster M, Ling Q, Schottler MA, Walther D, Zoschke R, Grimm B, Jarvis RP, Bock R (2019) Control of retrograde signalling by protein import and cytosolic folding stress. Nat Plants 5: 525–538

75. Zeng J, Geng X, Zhao Z, Zhou W (2024) Tipping the balance: The dynamics of stem cell maintenance and stress responses in plant meristems. Curr Opin Plant Biol 78: 102510

76. Zhang Y, He P, Ma X, Yang Z, Pang C, Yu J, Wang G, Friml J, Xiao G (2019) Auxin-mediated statolith production for root gravitropism. New Phytol 224: 761–774

77. Zhao Z, Andersen SU, Ljung K, Dolezal K, Miotk A, Schultheiss SJ, Lohmann JU (2010) Hormonal control of the shoot stem-cell niche. Nature 465: 1089–1092

